# Exploring the Role of Cotton *CMF* Genes in Salt Stress Tolerance: Insights from Phylogenetic, Expression, and Functional Analyses

**DOI:** 10.1101/2024.10.16.618649

**Authors:** Yupeng Cui, Gongyao Shi, Shuai Wang, Xingyu Peng, Xue-Rong Zhou, Wuwei Ye

## Abstract

Salt stress significantly limits cotton growth and development, making the improvement of salt tolerance critical. *CMF* genes, containing a single CCT domain, are promising targets for breeding programs. We identified 101 *CMF* genes from four cotton species. Phylogenetic analysis grouped these genes into six clusters closely related to *Arabidopsis*. Gene structure and motif analysis showed high conservation, while collinear analysis indicated segmental duplication as a key factor in *CMF* gene family expansion. Promoter analysis predicted involvement in light, abiotic stress, plant hormone responses, and growth. Expression analysis revealed differential expression of *GhCMF* genes across tissues, with some induced by abiotic stress. Co-expression network analysis indicated *GhCMF14* interacts with other proteins to resist abiotic stress. Subcellular localization showed GhCMF14 in the nucleus. Silencing *GhCMF14* enhanced salt tolerance, with higher chlorophyll and proline content, and lower MDA and H_2_O_2_ levels. This study provides insights into *CMF* gene regulation and their role in cotton salt stress tolerance.

## 1. Introduction

The CCT gene family can be divided into three subfamilies based on the conserved CCT domain [1]. These three subfamilies are CONSTANS-like (COL), pseudo-response regulator (PRR), and CCT motif family (CMF). COL subfamily proteins harbor one CCT domain and one or two additional zinc finger B-box (BBOX) domains, PRR subfamily proteins contain a CCT domain in the C-terminal and a pseudo-receiver domain in the N-terminal. CMF subfamily proteins do not have any characterized domains other than the CCT domain [2]. The CCT domain, a highly conserved 43-amino acid module, which located towards the C-terminal of proteins, and is involved in nuclear localization and protein-protein interactions [3, 4]. The proteins encoded by the CMF gene family have a variety of activities, however, the specific functions of these genes need further research to elaborate. Phylogenetic analysis showed that the BBOX domain originally present in the COL gene was gradually lost in the CMF gene lineage as evolution progressed in grasses, resulting in the CMF gene eventually retaining only one CCT domain [2].

The *CCT* gene family members not only participate in the circadian rhythm and development regulation of plants, but also may play an important role in coping with abiotic stress responses. For example, *AtCOL3* as a positive regulator of photomorphogenesis that promotes lateral root development and also acts as a daylength-sensitive regulator of shoot branching [5]. *AtCOL7* has a dual role, which could promote branching under high R:FR conditions and enhance shade avoidance syndrome under low R:FR conditions [6]. *Ghd2*, as a negative transcription factor in rice, is involved in drought-induced leaf senescence [7]. *AtCOL4* in *Arabidopsis* is involved in ABA and salt stress responses through ABA-dependent signaling pathways [8]. In *Tamarix hispida*, overexpression of *ThCOL2* can enhance ROS clearance to effectively reduce cell damage and death caused by salt stress [9]. In mango (*Mangifera indica* L.), *MiCOL1A* and *MiCOL1B* are involved in the flowering pathway and enhanced drought tolerance in *Arabidopsis* [10]. In apple (*Malus domestica*), Zinc-finger protein B-BOX 7/CONSTANS-LIKE 9 (MdBBX7/MdCOL9) formed a module with MdMIEL1, and played an active role in drought resistance in apples [11]. GmCOL1a is a nuclear localization protein that regulates salt tolerance and drought tolerance in soybean [12].

The *CMF* gene is a subfamily of the *CCT* gene family, which has been less studied than the other two subfamilies (*COL* and *PRR* subfamilies). Current studies have found that HEADING DATE 7 (*Ghd7*), one of the *CMF* subfamily member in rice, is involved in the growth and development of rice and the regulation of abiotic stress processes as a negative regulatory factor. Overexpression of *Ghd7* increases drought sensitivity, while knock-down of *Ghd7* enhances drought resistance [13]. In addition, *Ghd7* has been shown to regulate flowering, plant growth, and hormone metabolism [14, 15]. In maize (*Zea mays*), insertion of a CACTA-like transposable element into the *ZmCC* (a *CMF* gene) promoter could inhibiting *ZmCC* expression, and attenuate the sensitivity of maize under long-day conditions, which can lead to a sharp reduction in flowering time and facilitate the spread of maize cultivation to long-day environments [15]. Moreover, *ZmCCT* has dual functions in regulating flowering time and stress response, which negatively regulates flowering time and enhances drought resistance of maize under LD conditions [16, 17]. *FITNESS* (*AtCMF3*) plays a key role in regulating reactive oxygen species (ROS) level and defense responses [18]. *CIA2* (*AtCMF14*) and *CIL* (*AtCMF9*) are both important regulators of plant development, in which *CIA2* is the main regulator of chloroplast and flower organ development, which increases flower number and inflorescence axis height, *CIL* promotes the flowering process and mainly controls the timing of flowering [19]. It is worth noting that *AtCIA2*, *AtCIA2-like* and their homologs in albostrians *Barley* (*HvCMF3* and *HvCMF7*) are play an important role in chloroplast biogenesis, photosynthesis and abiotic stress responses (UV-AB, bright light and heat shock) [20–22]. The expression of 19 *CMF* genes, identified in soybean from transcription data, suggests that they are functionally diverse in metabolism and stress responses [23]. Twenty-five *CMF* genes have been identified in *Brassica rapa*, and *BrCMF5*, *BrCMF7*, *BrCMF14*, *BrCMF21* and *BrCMF22* play important roles in coping with abiotic stress [24]. Those research showed that different *CMF* members may have different biological functions.

Plants are exposed to a variety of external stimuli that affect the transition from the vegetative stage to the reproductive stage, as well as multiple processes such as flowering [25]. For example, when plants are subjected to salt stress, they affect seed germination, crop growth, and yield [26]. Cotton is an important cash and oil crop in China. Cotton provides 35% of the world’s total fiber usage, and its seeds are rich in fatty acids and proteins, which are widely used in processed edible oils, industrial raw materials and biodiesel [27]. However, stress conditions often affect the growth and development of cotton, thus reducing its yield. Different abiotic stresses may have different effects on cotton [28]. Currently, saline-alkali soils cover more than 800 million hectares, accounting for about 6% of the world’s total soil area [29]. Therefore, it is improtant to understand the genetic and regulation mechanism of salt tolerance in cotton and breed cotton varieties with excellent salt tolerance.

The identification and analysis of *CMF* genes have been reported in several species, including *Arabidopsis* [2, 30], rice [31], wheat [25], maize [32], poplar [33], soybean [23], etc. Moreover, the comprehensive analysis of *CONSTANS-like* genes in cotton has also been reported [34–37]. Although the *CMF* gene family has been reported to play important roles in plant development and stress tolerance, extensive characterization of cotton *CMF* genes has not yet been reported before. In this study, all of the *CMF* family genes in four cotton species were identified and characterized. The identified *CMFs* were comprehensively analyzed for phylogenetic classifications, gene structures, protein motifs, chromosomal locations, duplicated genes and cis-acting elements. Subsequently, the expression profiles of *GhCMF* genes were analyzed in different tissues and abiotic stresses. In addition, a representative gene *GhCMF14* that was differentially expressed under 300 mM NaCl treatment. Virus-induced gene silencing (VIGS) technology was used to preliminarily verify its function in response to salt stress. Our results provided a systematic view on the evolution and function of *CMF* genes in cotton.

## 2. Material and methods

### 2.1. Sequence retrial and identification of CMF family members in four cotton species

The genome and protein sequences of *Gossypium raimondii* (JGI) [38], *G. arboreum* (CRI) [39], *G. hirsutum* (ZJU) and *G. barbadense* (ZJU) [40] were obtained from Cotton functional genomic database (https://cottonfgd.org/) [41]. The published CMF proteins of rice and *Arabidopsis* [2, 23, 31] were sourced from the rice genome database (http://rice.plantbiology.msu.edu//) and the TAIR (https://www.arabidopsis.org/index.jsp) database, respectively. Afterwards, the BlastP program (E-value < 1×10−5) was performed to search for the candidate *CMF* genes of the four *Gossypium* species using AtCMF and OsCMF protein sequencesas queries. Then, each cotton CMF protein was subjected to the Pfam (http://pfam.sanger.ac.uk/) [42] and SMART (http://smart.embl-heidelberg.de/) [43] databases to confirm the presence of the CCT domain (PF06203). We also examine the detail information of cotton *CMF* genes, including the genomic lengths, protein length, molecular weights (MWs), isoelectric points (pIs), Grand Average of Hydropathy, and other biophysical properties, from CottonFGD [41] (Table S1). Furthermore, the subcellular localization was predicted using the online CELLO v2.5 server [44].

### 2.2. Sequences alignments and phylogenetic construction

A total of 101 CMF proteins, including 16 from *G. raimondii*, 17 from *G. arboreum*, 34 from *G. hirsutum*, 34 from *G. barbadense*, 15 from *Arabidopsis*, and 19 from rice, were used for the phylogenetic analysis of CMF proteins in plants. The 135 CMF protein sequences were aligned with Clustal X v2.0 software [45] with default parameters, and followed by manual comparisons and refinements. Subsequently, we constructed the phylogenetic tree in MEGA 7.0 software under Neighbor-joining (NJ) method with pairwise deletion option, poisson correction model, uniform rates and bootstrapping with 1000 replications [46].

### 2.3. Analyses of gene structure and conserved motif

The exon-intron structures of cotton *CMF* genes were graphically visualized using the Gene Structure Display Server (http://gsds.gao-lab.org/) tool [47], based on the predicted *CMF* coding sequences and corresponding genomic sequences. The conserved motifs in cotton CMF proteins were analyzed using the MEME software (http://meme-suite.org/). The number of repetitions was set as any, the optimum width of motifs ranged from 10 to 50 residues, the maximum number of motifs was 3, and the other parameters remained at default values.

### 2.4. Chromosomal localization and gene duplication

The chromosome location images of 101 cotton *CMF* genes were mapped by Mapchart v2.2 software [48]. The gene duplication events of four cotton species were analyzed by using MCScanX software, and according to the length of aligned sequence, covered >80% between aligned gene sequences, similarly to the aligned regions (>80%) [49, 50]. Orthologous and homoeologs for *CMF* genes were identified with MCScanX software [51] and results were displayed using simple Circos-0.69 software [52].

### 2.5. Selective pressure analysis of duplicated gene pairs

To investigate the selection pressure experienced by *CMF* duplicated gene pairs from the four cotton species, all the full-length gene sequence of cotton *CMF* duplicated gene pairs of four cotton species were aligned by Clustal X v2.0 software firstly, then the synonymous substitution (Ks) and nonsynonymous substitution (Ka) were calculated using the DnaSP v5.0 software [53]. Finally, the Ka/Ks ratio was used to estimate the selection pressure for each gene pair.

### 2.6. Promoter cis-acting element analysis and expression patterns of GhCMF genes in cotton

To identify the cis-elements in the promoter sequences of the 34 *GhCMF* family genes, 2.0 kb promotor sequences of *GhCMFs* were extracted from CottonFGD database [41], and their regulation elements were predicted using the PlantCARE database (http://bioinformatics.psb.ugent.be/webtools/plantcare/html/). The identities and locations of the cis-elements of *CMF* promoters were visualized via TBtools [54]. The RNA-Seq data of *G. hirsutum acc* TM-1 was downloaded from the NCBI under the accession number SRA: PRJNA248163 [55] to analyze the expression level (FPKM) of *GhCMF* family genes at different developmental stages, and under cold, heat, salt and PEG stress. The heatmap was drawn by MultiExperiment Viewer (MeV) software based on the different expression levels of *GhCMF* family genes.

### 2.7. salt stress treatment and quantitative real-time PCR (qPCR)

Cotton seeds of *G. hirsutum* L. *acc* Zhong 9807 were obtained from the Institute of Cotton Research of the Chinese Academy of Agricultural Sciences. Cotton seedling of Zhong 9807 were placed in a plant growth chamber at 25 ± 2℃ under a 16 h light/ 8 h dark cycle. The third true leaf stage seedlings were treated with salt stress. Seedlings were grown in Hoagland nutrient solution and treated with additional 300 mM NaCl for 0, 12 and 24 h. The leaves were harvested at each time point of each treatment, and immediately frozen with liquid nitrogen, then stored at −80 ℃ for subsequent RNA isolation.

The RNAprep Pure Plant Kit (Tiangen, Beijing, China) and the PrimerScript 1st Strand cDNA Synthesis Kit (TaKaRa, Dalian, China) were used to extract total RNA from all samples and synthesis of first-strand cDNAs. Primer v5.0 software was used to design the gene-specific primers, according to the CDSs of GhCMF genes. The specific primers used are listed in Table S2. GhUBQ7 was used as an internal control to normalize all data. The qRT-PCR was strictly performed with SYBR premix Ex Taq Kit (TakaRa) using ABI 7500 real-time PCR System (Applied Biosystems, Foster City, CA, USA) based on the the manufacturer’s instruction. qRT-PCR was performed at 95 °C for 3 min, followed by 40 cycles of 95 °C for 5 s, 57 °C for 15 s, and 72 °C for 30 s in a 96-well plate. Three biological replicates and three technical replicates were conducted for each sample. The 2^-ΔΔCT^ method was used to calculate the relative expression levels of *GhCMF* genes [56].

### 2.8. Co-expression network analysis of GhCMF14

In order to examine the interaction network of GhCMF14 protein, we acquire ortholog gene by comparing the protein sequence of GhCMF14 to *Arabidopsis*. STRING software (https://string-db.org/) was used to analyze the interaction of GhCMF14 protein on the basis of the ortholog in Arabidopsis with a confidence parameter set at 0.4 threshold.

### 2.9. Subcellular location, virus-induced gene silencing (VIGS) and salt treatment

*GhCMF14* gene was amplified with gene specific primers from Zhong 9807. Using ClonExpress RII One Step Cloning Kit (Vazyme, Nanjing, China), the PCR products of *GhCMF14* genes was cloned into digested 2300-YFP vector with BamHI and SacI, respectively. Subcellular location process was carried out as previously reported [57]. YFP-GhCMF14 plasmid was transformed into *Agrobacterium* GV3101 by freeze-thaw method. The Agrobacterium culture carrying relative plasmid was infiltrated into 4-week old *Nicotiana bentamiana* (*N. bentamiana*) leaves. The plants were then maintained in a culture room with a temperature of 22℃ and a relative humidity of 60%, for 2 days, followed by observing fluorescence under a LSM780 confocal laser microscope with excitation at 514 nm for YFP.

The purified 300-bp fragment of *GhCMF14* was inserted into the pYL156 vector through the restriction sites BamHI and SacI. Recombinant plasmid pYL156:GhCMF14 was transformed into *Agrobacterium* GV3101 by freeze-thaw method. The empty vector pYL156 was employed as negative control. pYL156:PDS was as positive control. The VIGS procedure was carried out in accordance with our article previously described [57]. When the leaves of positive control plant turned white, indicating that the experiment was successful. After 3 weeks of infiltration, seedlings were treated with 300 mM NaCl solution for 0 and 24 h. The leaves were harvested at each time point of each treatment, and the silencing efficiency of *GhCMF14* was tested by qRT-PCR. The specific primers used are listed in Table S2.

### 2.10 Physilogical index measurement

After seedlings of VIGS plants were treated for 0 and 24 h under 300 mM NaCl, the chlorophyll contents of VIGS plants were measured by using a SPAD-502 type chlorophyll meter, which gave SPAD values. The leaf temperature and relative humidity remained at 24°C and 45%, respectively, and the photon flux density was 160 to 200 μmol m^-2^ s^-1^. Data were recorded after equilibration (approximately 6 min). The activities of superoxide dismutase (SOD) and peroxidase (POD), and the Content of MDA and Proline were determined by using kits purchased from Beijing Solarbio Science (China). All measurements were performed according to the manufacturer’s instructions.

### 2.11. DAB staining

The 3,3′-diaminobenzidine (DAB) working solution was made according with the instructions of enhanced DAB chromogenic kit, and kept chilled at 4°C. cotton leaves of VIGS plants were put into DAB dyeing solution, and dark incubate at room temperature for 24h. The leaves were removed from the dyeing solution and soaked in anhydrous ethanol until the green color of the leaves completely faded away. Then, photos were taken to observe the distribution and accumulation of ROS.

### 2.12. Statistical analysis

The results were analyzed by Student’s t-test. Differences were considered statistically significant when p < 0.05 (*) and p < 0.01 (**) compared with the control.

## 3. Results

### 3.1. Identification of CMF gene family in Gossypium

We used the protein sequences of 15 AtCMF and 19 OsCMF genes to search for related cotton CMFs in diploid cotton (*G. raimondii* and *G. arboreum*) and tetraploid cotton (*G. hirsutum* and *G. barbadense*) protein databases. Through comprehensive screening, a total of 101 cotton CMF sequences were identified, including 16, 17, 34 and 34 CMF members in *G. raimondii*, *G. arboreum*, *G. hirsutum* and *G. barbadense*, respectively. The number of *CMF* genes in each of the diploid cotton (*G. raimondii* and *G. arboreum*) exceeded that in *Arabidopsis* and rice, and *G. hirsutum* or *G. barbadense* contained the largest number of *CMF* genes, which were equal to the sum of that of the two diploid cotton species. These *CMF* genes were designated as *GrCMF1*-*GrCMF16*, *GaCMF1*-*GaCMF17*, *GhCMF1*-*GhCMF34* and *GbCMF1*-*GbCMF34* in the four cotton species, respectively, according to the ascending order of their chromosomal positions. Detailed information of all the CMF gene members was list in Table S1. Among the four cotton species, the length of 101 cotton *CMF* genes ranged from 405 bp (GaCMF16) to 1287 bp (*GhCMF2* and *GhCMF19*), and their corresponding proteins varied from 134 (GaCMF16) to 428 (GhCMF2 and GhCMF19) amino acids in length; the predicted molecular weight of the CMF proteins in the four cotton species ranged from 15.687-46.992 kDa, while the theoretical pI ranged from 4.197-10.255. The prediction of the subcellular localization of cotton CMF protein showed that 100 proteins are basically located in the nuclear, while GaCMF16 was in the nuclear or mitochondria (Table S1).

### 3.2. Phylogenetic relationship analysis in cotton CMF genes

To evaluate the evolutionary relationships of the 135 CMF proteins among cotton (G*. raimondii*, *G. arboreum*, *G. hirsutum* and *G. barbadense*), *Arabidopsis* and rice, we constructed a phylogenetic tree by MEGA 7.0 using a NJ method. The results show that the 135 CMF family members from six species were separated into six groups (Group I, Group II, Group III, Group IV, Group V, and Group VI) based on the phylogenetic tree (Fig. 1). And the member numbers of each group were compared among these six plant types (Fig. S1). We found that group V had the highest numbers of *CMF* family genes (28), followed by Group I (26) and Group VI (26), Group IV (22), Group II (21), and Group III (12) among the six different plant species. Interestingly, *OsCMF* family genes have no member in Group I and III, which suggesting that the members of the two groups among the five different plant species are relatively less close to members of Groups II, IV, V, and VI. In addition, the cotton *CMF* genes showed a closer relationship to the *AtCMF* genes than the *OsCMF* genes, which is consistent with our understanding of the evolutionary history of these genes in plants. The number of *CMF* genes in *G. hirsutum* or *G. barbadense* was twice of that in *G. raimondii* or *G. arboreum*, and most *CMF* genes in diploid cotton corresponds to two orthologous genes in tetraploid, since approximately 1-2 million years ago, an A-genome like ancestral African species, *G. herbaceum* or *G. arboreum* and a native D-genome-like species, *G. raimondii* (D5) hybridized and underwent polyploidization, giving rise to *G. hirsutum* and *G. barbadense* [40, 55, 58]. The inconsistencies were that there were no corresponding *CMF* genes in *G. arboreum* for the two genes (*GhCMF1* and *GbCMF1*) from A subgenome of *G. hirsutum* and *G. barbadense*, and also no *CMF* genes in *G. raimondii* for the two genes (*GhCMF25* and *GbCMF25*) from D subgenome of *G. hirsutum* and *G. barbadense*, suggesting that the two genes were lost after *G. arboreum*/*G. raimondii* differentiation.

**Fig. 1.**
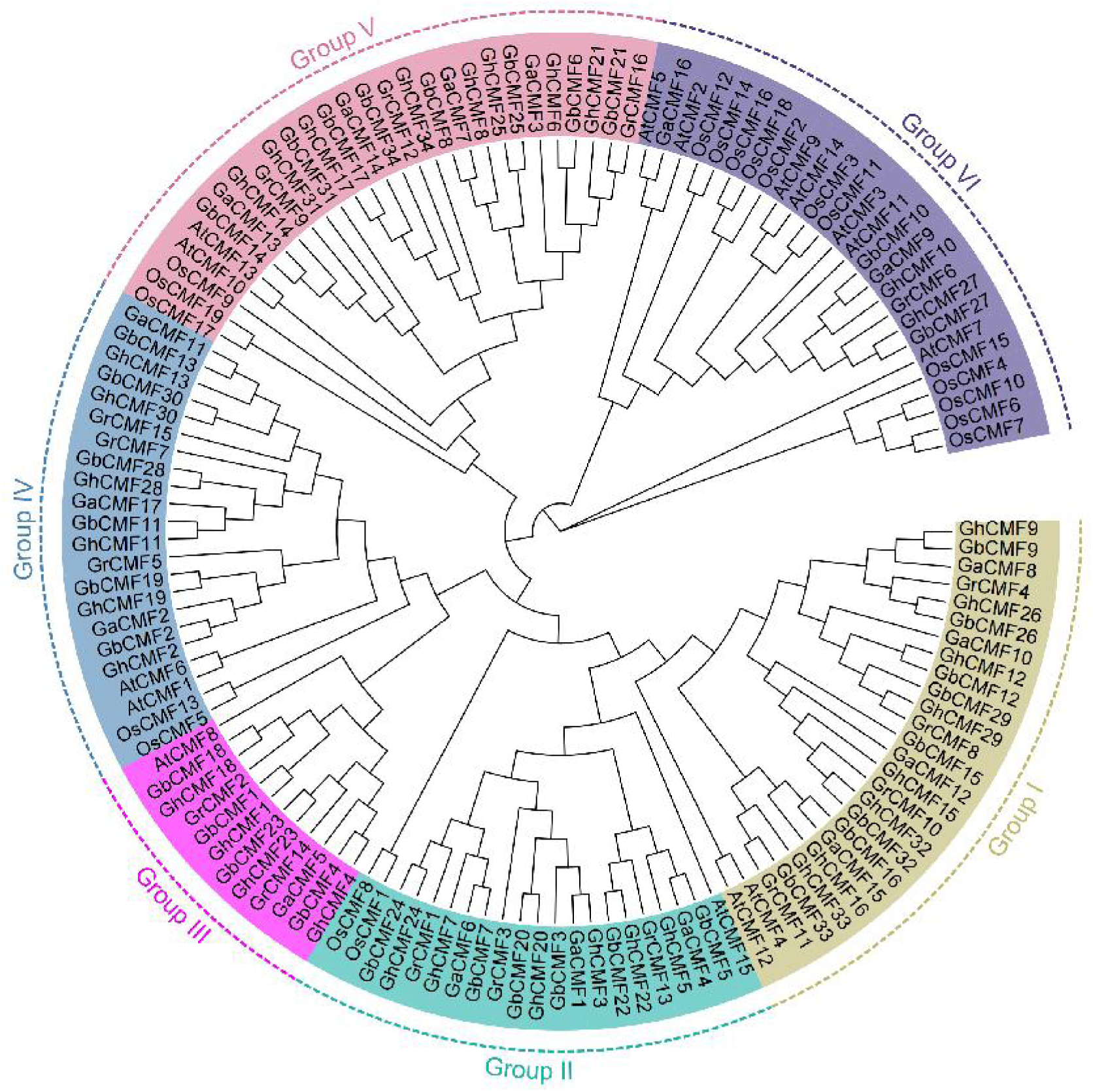
Phylogenetic analysis of the CMF protein family. Gr, *G. raimondii*; Ga, *G. arboretum*; Gh, *G. hirsutum*; Gb, *G. barbadense*; At, *Arabidopsis thaliana*; Os, *oryza sativa*.

### 3.3. Gene structure, conserved motifs and chromosome mapping analysis of the cotton CMF genes

Differences in gene structure play an important role in gene evolution [59]. Further information on conservation and diversity of cotton *CMF* genes was obtained by analyzing its gene structure. As shown in Fig. S2A, the structural analysis revealed a considerable variation in the length and structure of cotton *CMF* family genes. Among the 101 cotton *CMF* genes, *GaCMF16* was the smallest gene (405 bp), while *GhCMF2* and *GhCMF19* were the largest gene (1287 bp). The cotton *CMF* genes contained multiple exons ranging from two to six. Although the number of exons varied among the six groups, the exon patterns within groups were similar. For example, the majority of *CMF* genes in Group I contained four (22/24) exons, except for *GrCMF11* and *GaCMF15*, which contained 5 and 6 exons, respectively. All members in Group VI had three exons except *GaCMF16*, which contained two exons. In addition, we observed differences in the number of introns among the *CMF* genes in the four cotton species, and the variation of the total length and number of introns mainly was contributed to the wide-ranging sizes of cotton *CMFs* genomic sequence, and the same phenomenon could be seen in other gene families [60, 61].

Conserved motifs of the four cotton CMF proteins were also analyzed using the MEME software. The motif length and CMF motif number were 50 AA and 1-3, respectively. A total of 3 motifs were identified (Fig. S2B), and the details of each motif were shown in Fig. S3. Motifs 1was annotated as the CCT motif, which was widely presented among all cotton CMF protein. In the four cotton species, several residues at positions 14, 17, 21, 23, 27, 29, 32, 33, 36, 41, 44, 45, and 46 of the CCT motif consensus sequence were identical, and the rest of them were also highly conserved (Fig. S3). Moreover, we can found that proteins of the same group usually shared group-specific conserved motifs (Fig. S2B), which was similar to the evolutionary tree and gene structure. For example, motif 2 and motif 3 was exclusively present in the members of Group V except GaCMF2 in Group IV, which suggested that they may have special functions.

In order to understand the specific distribution of genes on chromosomes more intuitively, we constructed chromosome distribution maps of the cotton *CMF* genes in the four cotton species. As shown in Fig. 2, 100 of 101 cotton *CMF* genes were unevenly assigned to chromosomes, and only one was not anchored on chromosome. In *G. raimondii*, 16 *GrCMF* genes were widely distributed on 11 chromosomes. Three genes were located on chromosome 6, followed by two on chromosomes 7, 8, and 9, and the remaining chromosomes had a single member. In *G. arboreum*, except for *GaCMF17*, the remaining16 *GaCMF* genes were distributed on 11 of 13 chromosomes. Among them, two members were distributed on chromosomes 5, 8, 9, 11, and 12, and only one member on the remaining chromosomes. All *GhCMF* genes were dispersed across 22 of the 26 *G. hirsutum* chromosomes, except for A06, A13, D06 and D13. Two chromosomes (A09 and D09) each contained three genes, eight chromosomes (A05, A08, A11, A12, D05, D08, D11 and D12) each contained two genes, and the only one member was located on the remaining chromosomes. The gene localization of *G. barbadense* is the same as that of *G. hirsutum*.

**Fig. 2.**
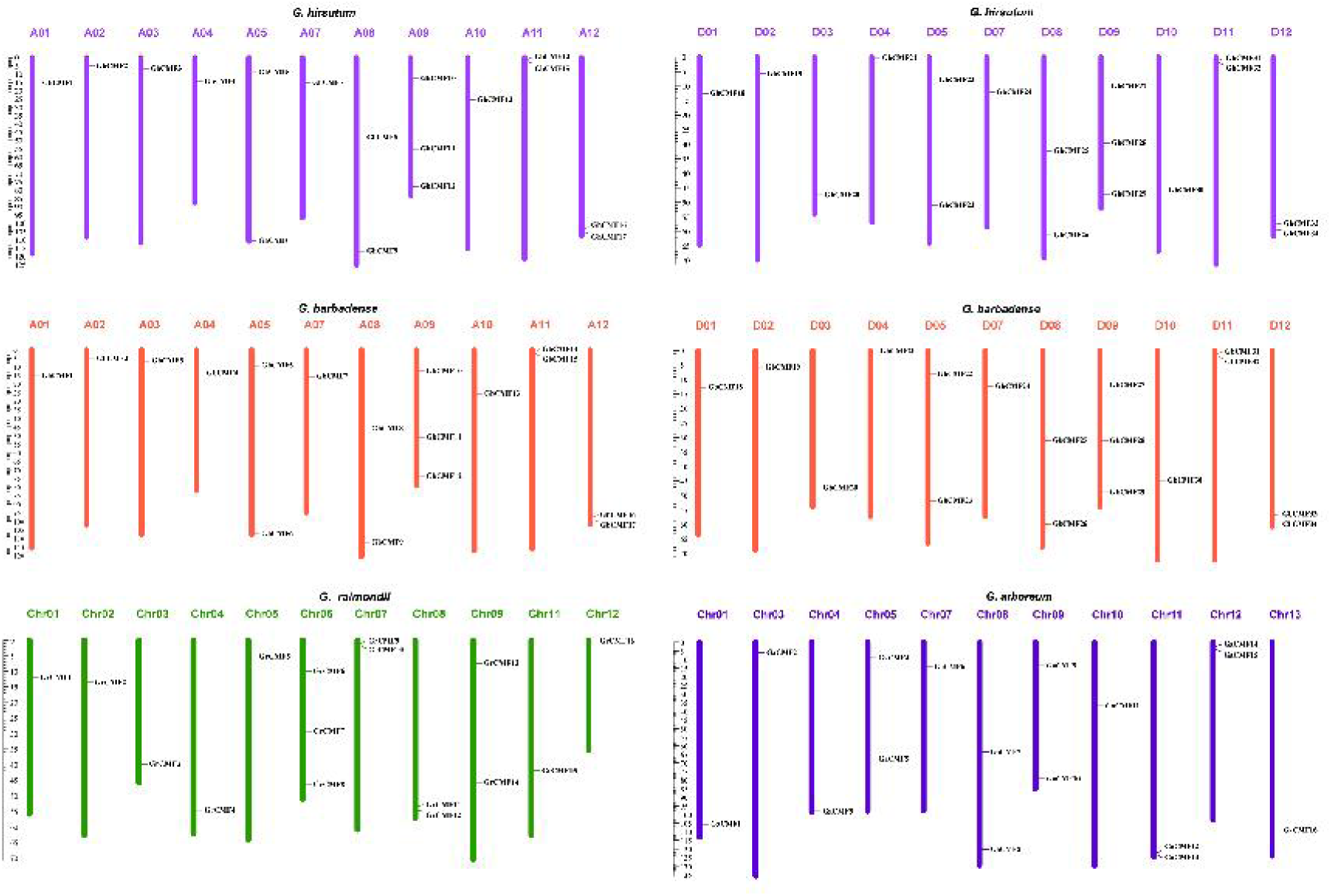
Chromosome distribution of *CMF* gene families in four cotton species.

### 3.4. Gene duplication and selection pressure analysis in cotton CMF family genes

Gene duplication is a mechanism for the rapid generation of additional sequences for natural selection to act and confer greater organismic fitness [62]. To better study the driving force for the evolution and the functional divergence of cotton *CMFs*, we further explored gene duplication events. As shown in Fig. 3 and Table S3, a total of 410 pairs of homologous genes exhibited a collinear relationship in the four cotton species. Among them, 11, 16, 85, and 91 pairs of duplicated *CMF* genes were identified in *G. raimondii*, *G. arboreum*, *G. hirsutum* and *G. barbadense*, respectively (Fig. S4). And 207 duplicated *CMF* genes were identified from inter genomic combinations of the four cotton species (Fig. S5). All *CMF* duplicated gene pairs were involved in segmental duplication because they were located on individual chromosomes or the distance between these genes on the same chromosome were more than 100 kb. This suggest that segmental duplication events were mainly contributed to the expansion of the *CMF* family genes in the four cotton species.

**Fig. 3.**
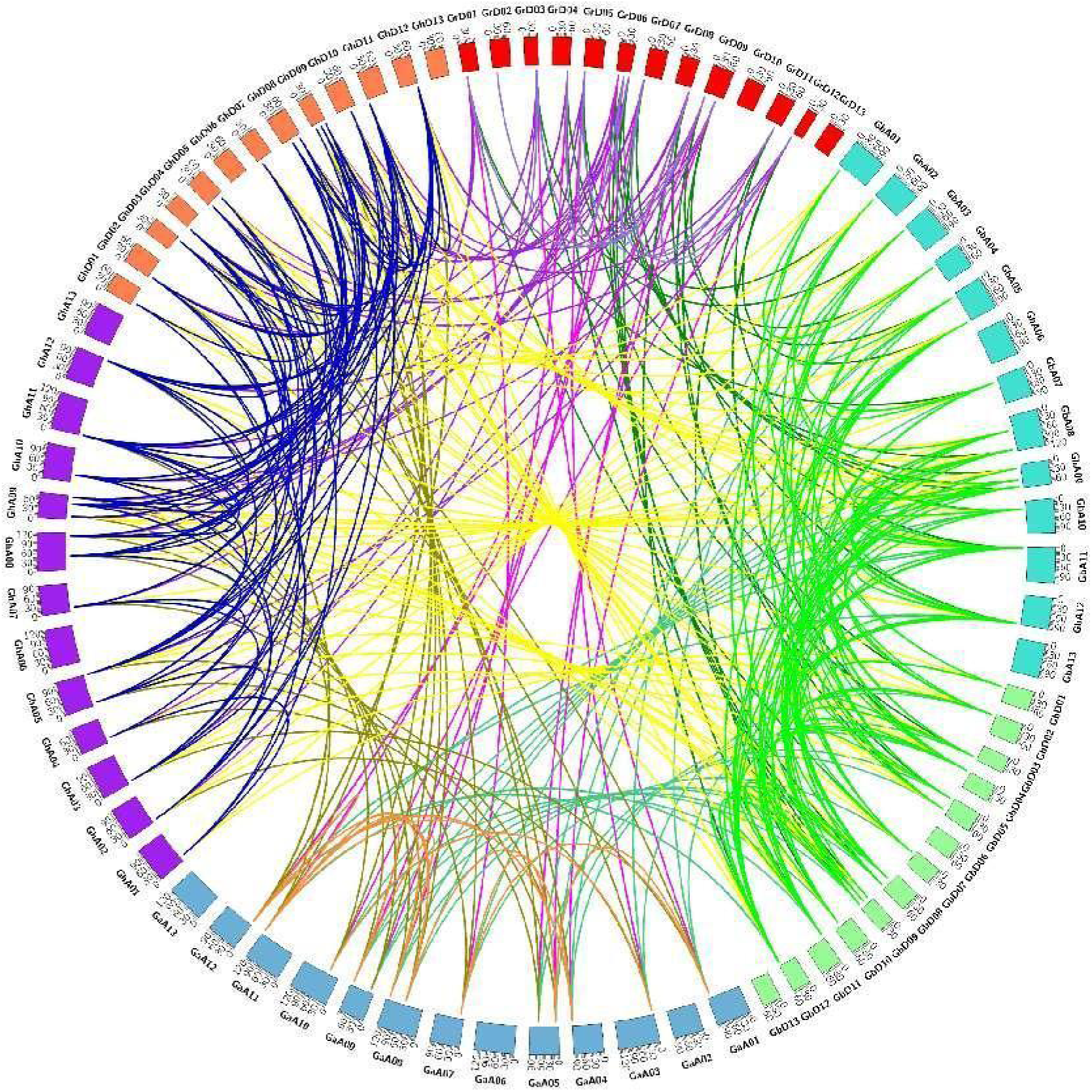
Syntenic relationship of *CMF* duplicated genes pairs from four cotton species (*G. hirsutum*, *G. barbadense*, *G. arboreum* and *G. raimondii*).

During the course of evolution, duplicated gene pairs might have undergone three alternative fates, i.e., nonfunctionalization, subfunctionalization and neofunctionalization [63, 64]. In this study, we calculated the Ka/Ks ratio of 410 cotton *CMF* gene duplicated pairs of the four cotton species (Table S4), which aim to explore the selective constraints on their divergence after duplication. It showed that 400 duplicated gene pairs with a Ka/Ks ratio below one, which experienced strong purifying selection, while 10 duplicated gene pairs were greater than one (positive selection), illustrating the relaxed selection pressure during evolution. Among them, 352 pairs of duplicated *CMFs* with a Ka/Ks ratio below 0.5 were found from inter and intra genomic of four cotton species, and 48 pairs were ranging from o.5 to 1 (Table S4), suggesting that synonymous substitutions occurred more frequently than non-synonymous ones, and the functional constraint with strong purifying selection during evolution. Moreover, it could be found 156 of 207 orthologue pairs with Ka/Ks ratio below 0.5 from inter genomic of four cotton species, indicating those gene pairs had experienced strong purifying selection pressure after segmental duplication, which make their functions tend to be relatively similar. In additon, the duplication time period for cotton *CMF* genes were further calculated as shown in Table S4. The segmental duplication of *CMFs* in *G. raimondii* and *G. arboreum* occurred from 14.96 Mya to 33.54 Mya and 13.80 Mya to 33.90 Mya, respectively. While, the *CMF* duplication gene pairs occurred from 0.10 Mya to 38.12 Mya and 0.10 Mya to 37.31 Mya in *G. hirsutum* and *G. barbadense*, respectively.

### 3.5. Anaktsis of cis-acting elements in GhCMF Promoter, and GhCMF genes with altered expression

Globally, more than 90% of cultivated cotton species is upland cotton due to its higher fiber yield and environmental adaptability [65]. And analyzing the cis-acting elements in genes can provide a lot of important evidence for understanding the function of genes. To identify conserved cy-elements present in the proximal promoter region of *GhCMF* gene, bioinformatic analysis of 34 upstream 2kb promoter regions of *GhCMF* gene was performed using PlantCARE online software. Besides some common and core cis-elements, such as the CAAT-box and the TATA-box, nine patterns of cis-element motifs were identified upstream of the 34 *GhCMF* genes (Fig. 4A, Table S5). These cis-elements were further categorized into four classes: light responsiveness, abiotic stress responsiveness, plant hormone responsiveness, and growth regulation [66]. The analysis showed that all 34 *GhCMF* promoter regions contained light-responsive elements, while only six *GhCMF* promoter regions contained growth regulatory elements, suggesting that *GhCMFs* may be involved in stress and light responses. Abiotic stress-responsive elements include four types of elements: anaerobic induction responsive element, drought-inducibility responsive element, defense and stress responsive element, and low-temperature responsive element. Twenty-eight *GhCMF* promoter regions contained anaerobic induction responsive element, 13 contained drought-inducibility response elements, 14 contained defense and stress responsive element, and 11 contained low-temperature response elements. Plant hormone response elements include three different types of elements (abscisic acid responsive element, gibberellin responsive element, and MeJA responsive element). 21 *GhCMF* promoter regions contained abscisic acid responsive elements, 15 contained gibberellin response elements, and 25 contained abscisic acid responsive elements (Table S6). The results suggested that *GhCMFs* with these cis-acting elements may be involved in a variety of regulatory mechanisms in cotton under different stress conditions.

**Fig. 4.**
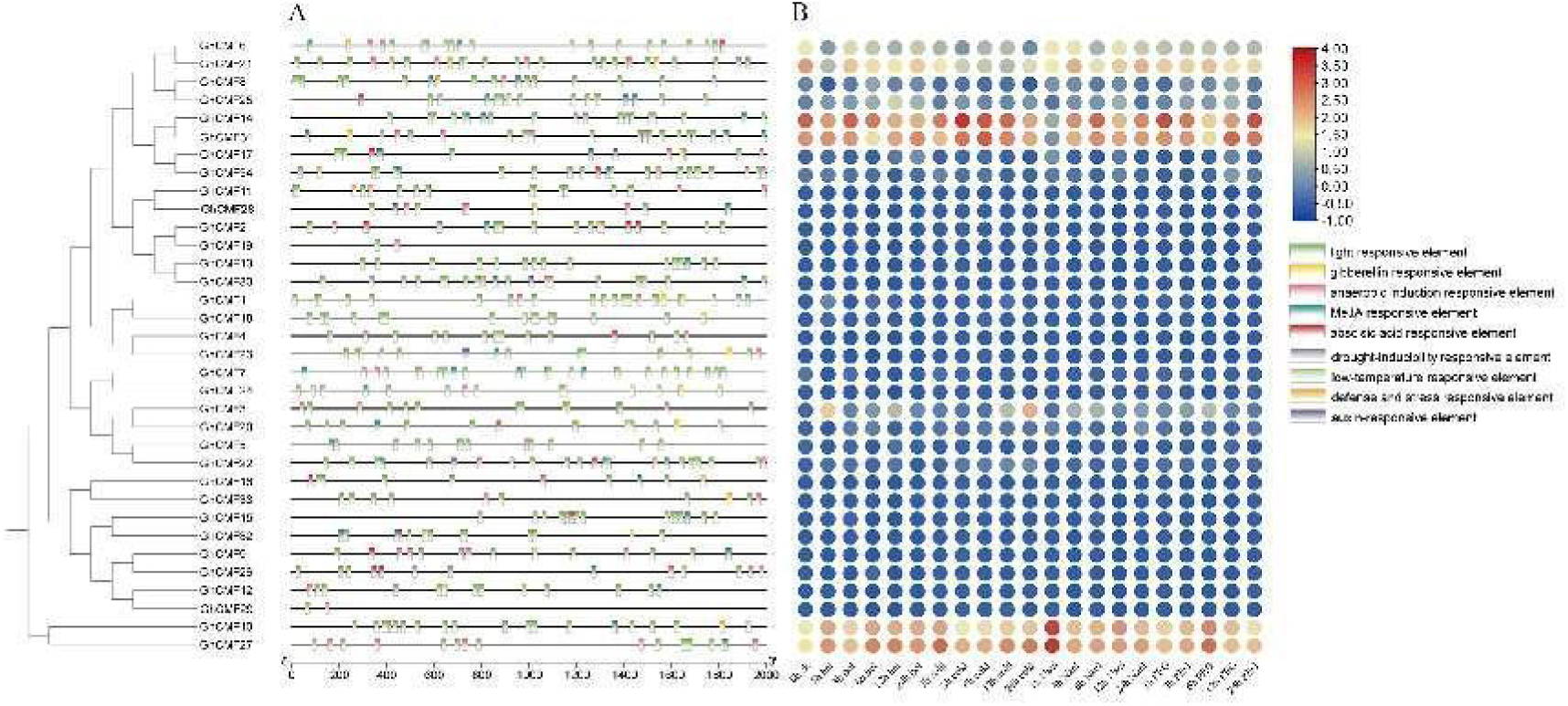
Expression patterns and promoter analysis of the *GhCMFs*. (A) Cis-acting elements in promoters of *GhCMFs*. (B) Heatmap of the expression of *GhCMFs* under different abiotic stresses at diferent times of stress (hot, cold, salt and PEG).

To further investigate the response mechanisms of *GhCMF* genes to abiotic stress, transcriptome data of leaves treated with various abiotic stresses, including heat, cold, salt, and PEG, were retrieved (Fig. 4B). *GhCMF* genes participated in the control of abiotic stress, and genes within the same subfamily showed similar expression patterns. Under high-temperature stress, five *GhCMF* genes were highly upregulated in leaves, namely *GhCMF10*, *GhCMF21*, *GhCMF25*, *GhCMF27*, and *GhCMF31*, while *GhCMF3*, *GhCMF17*, and *GhCMF20* were weakly upregulated. Under cold stress, three *GhCMF* genes were significantly upregulated in leaves (*GhCMF3*, *GhCMF10*, and *GhCMF27*), and seven were downregulated (*GhCMF6*, *GhCMF8*, *GhCMF14*, *GhCMF17*, *GhCMF21*, *GhCMF31*, and *GhCMF34*). Under salt stress, *GhCMF10* and *GhCMF27* were highly upregulated in leaves, with *GhCMF3* and *GhCMF20* weakly upregulated. *GhCMF6*, *GhCMF14*, *GhCMF21*, and *GhCMF31* exhibited significantly downregulated. Interestingly, the expression of *GhCMF14* gene was significantly upregulated after 6h of salt stress, which was the most upregulated gene among all *GhCMF* genes. Under PEG stress, *GhCMF14*, *GhCMF31*, *GhCMF27*, and *GhCMF10* genes were significantly upregulated in leaves, while *GhCMF6* and *GhCMF21* were significantly downregulated. All these *GhCMF* genes showed significant upregulated or downregulated, suggesting their potential role in the cotton plant’s response to abiotic stress. Moreover, except for the duplicated gene pair *GhCMF3* and *GhCMF20*, which exhibited different expression patterns under high, low, salt, and PEG stress, other duplicated gene pairs, especially *GhCMF6/GhCMF21*, *GhCMF8/GhCMF25*, *GhCMF14/GhCMF31*, *GhCMF17/GhCMF34*, *GhCMF5/GhCMF22*, and *GhCMF10/GhCMF27*, showed similar expression patterns, which may imply that these duplicated genes evolved toward conservative functional evolution when responding to abiotic stress.

The expression levels of the *GhCMF* gene family members in roots, stems, and leaves were analyzed (Fig. 5A). In leaves, one duplicated gene pair, *GhCMF10*/*GhCMF27*, exhibited the highest expression levels. In roots and stems, one duplicated gene pair, *GhCMF14/GhCMF31* exhibited the highest expression levels. Additionally, 10, 11, and 8 genes showed no expression in leaves, roots, and stems respectively, and five genes (*GhCMF2*, *GhCMF19*, *GhCMF28*, *GhCMF11*, *GhCMF30*) were not expressed in any of these tissues (Fig. 5A, Table S7). The results demonstrated that the *GhCMF* gene family exhibits differential expression across different tissues, primarily associated with their functional roles in these tissues.

**Fig. 5.**
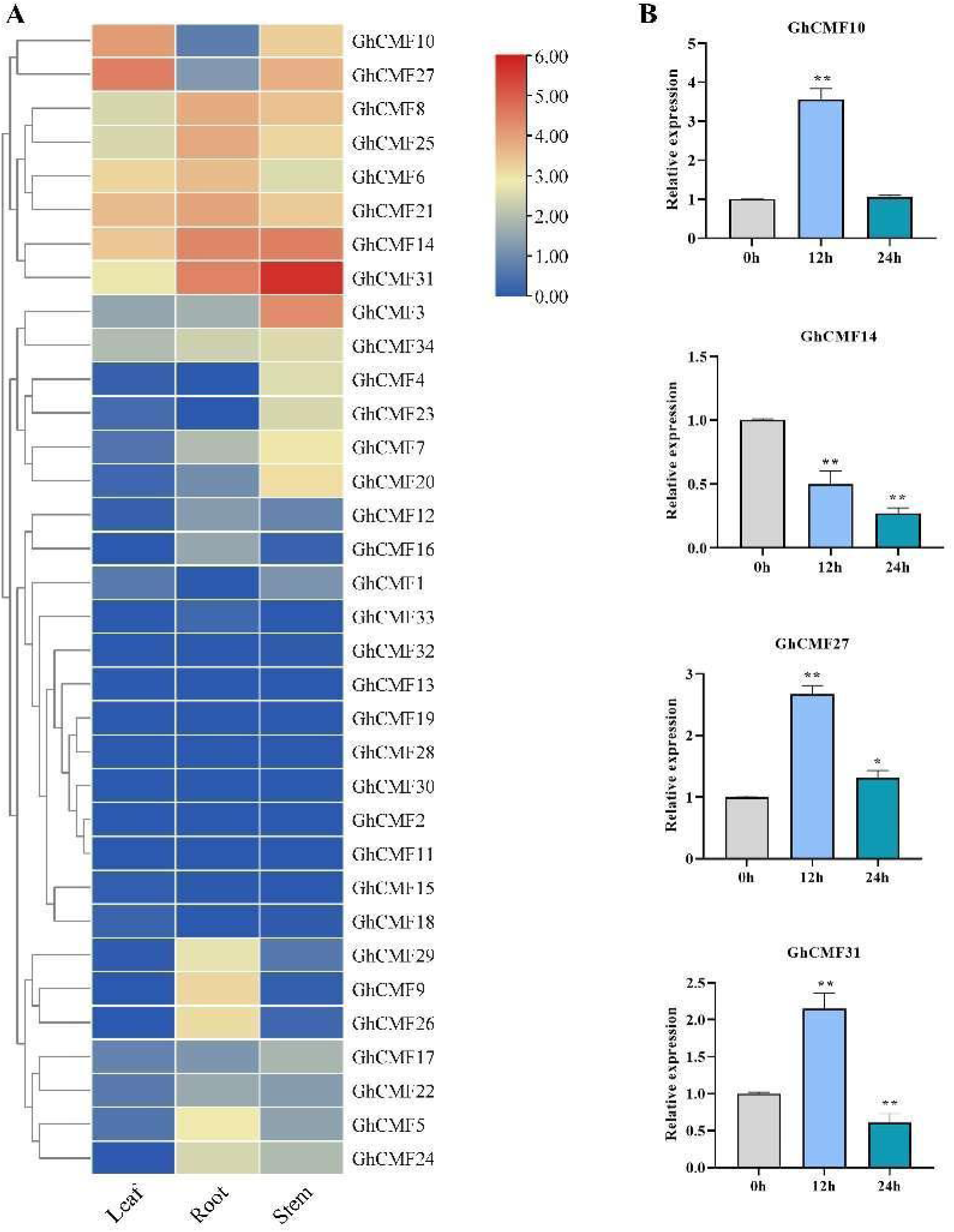
Heatmap of expression profiles of GhCMF family genes in different tissues (A) and relative expression of four *GhCMF* genes leaves of Zhong 9807 under NaCl treatment. Columns and bars separately represent the means and standard deviation (n= 3), and the data was determined by the t-test, Asterisks indicate significant differences compared with the 0 h (* p < 0.05, ** p < 0.01).

### 3.6. Investigation of GhCMF genes expression under salt stress

To further confirm the expression pattern of *GhCMF* gene in the transcriptome data under salt stress, we selected four *GhCMF* genes to examined their expression under salt stress in Zhong9807 material. The results showed that all four selected genes could be stimulated by salt stress (Fig. 5B). At 12h following exposure to salt treatment, the expression level of *GhCMF10* increased significantly at 12h of salt stress, but returned to normal at 24 h. *GhCMF14* decreased significantly at 12 h and further decreased at 24 h with prolonged salt treatment time. The expression level of *GhCMF27* was significantly up-regulated under salt treatment, and reached its peak at 12h. The expression level of *GhCMF31* was significantly increased at 12h, but significantly decreased at 24 h. These results suggest that these four *GhCMF* genes may play important roles in the response to salt stress in cotton.

### 3.7. Interaction network of GhCMF proteins

To further investigate the role of GhCMF protein, combined with promoter analysis and stress expression analysis, we selected *GhCMF14* as the gene for further research. The GhCMF14 protein sequence had the highest similarity with AT5G53420 based on the protein sequence search and comparison using STRING software, and 20 proteins were predicted (Fig. 6). Among them, 10 proteins (ERCC1, CR88, ZML2, ZML1, PRR7, ARP6, BBX15, BBX14, BBX16, and AT1G49390) were interacts with GhCMF14 (Figure S5B). Among them, BBX proteins form a subgroup of zinc finger transcription [67], consisting of 32 members in *Arabidopsis* [68]. *BBX* transcription factors play a critical role in a variety of life activities, including photomorphogenesis in different plant species [69], flowering [70], and biological and abiotic stress responses [71]. Moreover, recent studies have found that *BBX14* is a negative regulator of nitrogen starvation and dark-induced leaf senescence [72], and heterologous expression of *CpBBX19* in *Arabidopsis* could enhance the tolerance transgenic plant to salt and drought stress [71]. *ERCC1* was involved in the repair of oxidative damage, and increased *ERCC1* expression could compensate for increased oxidative stress due to decrease in photosynthetic electron sink [73]. *CR88* gene encodes a *HSP* gene, plays a very important role in normal development, heat stress and salt stress [74, 75]. ACTIN-RELATED PROTEIN 6 (ARP6), an essential subunit of SWR1-C, is implicated in flowering and stress response, ectopic expression of *AcARP6* in *Arabidopsis* delayed flowering and increased tolerance to salinity [73], and *Arabidopsis arp6* mutant was more sensitive to salt stress than the wild type [76]. ARP6 and SWC6 could interact with phyB, ARP6 and phyB could co-regulate numerouus genes in the same direction, some of which were associated with auxin biosynthesis and response including YUC9 [77]. This indicated that GhCMF14 may interact mutually to participate in cotton abiotic stress, especially salt stress.

**Fig. 6.**
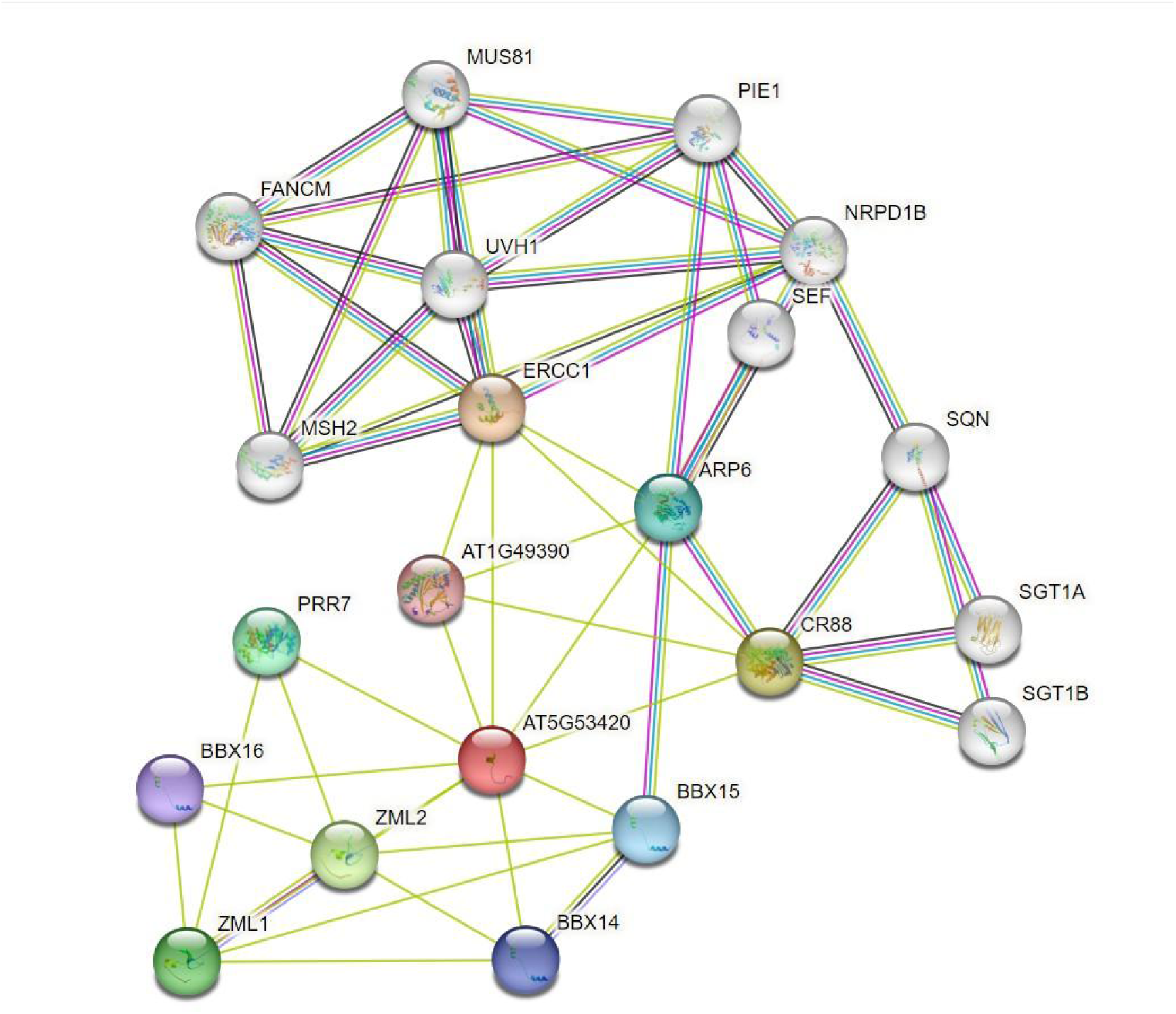
Interaction network of GhCMF14 protein.

### 3.8. Subcellular localization of GhCMF14

The tobacco transient transformation method was used to initially determine the distribution of GhCMF14 protein in cells to further understand its function. The *N. bentamiana* mesophyll cells were transformed transiently using the constructed vectors of YFP-GhCMF14, and the fluorescence signal was observed by laser scanning confocal microscope. The result showed that the YFP fluorescence signal was scattered throughout the cell, including the nucleus, whereas YFP-GhCMF14 was observed in the nucleus and the signals overlapped with H2B in the nucleus, indicating that GhCMF14 was localized exclusively in the nucleus (Fig. 7).

**Fig. 7.**
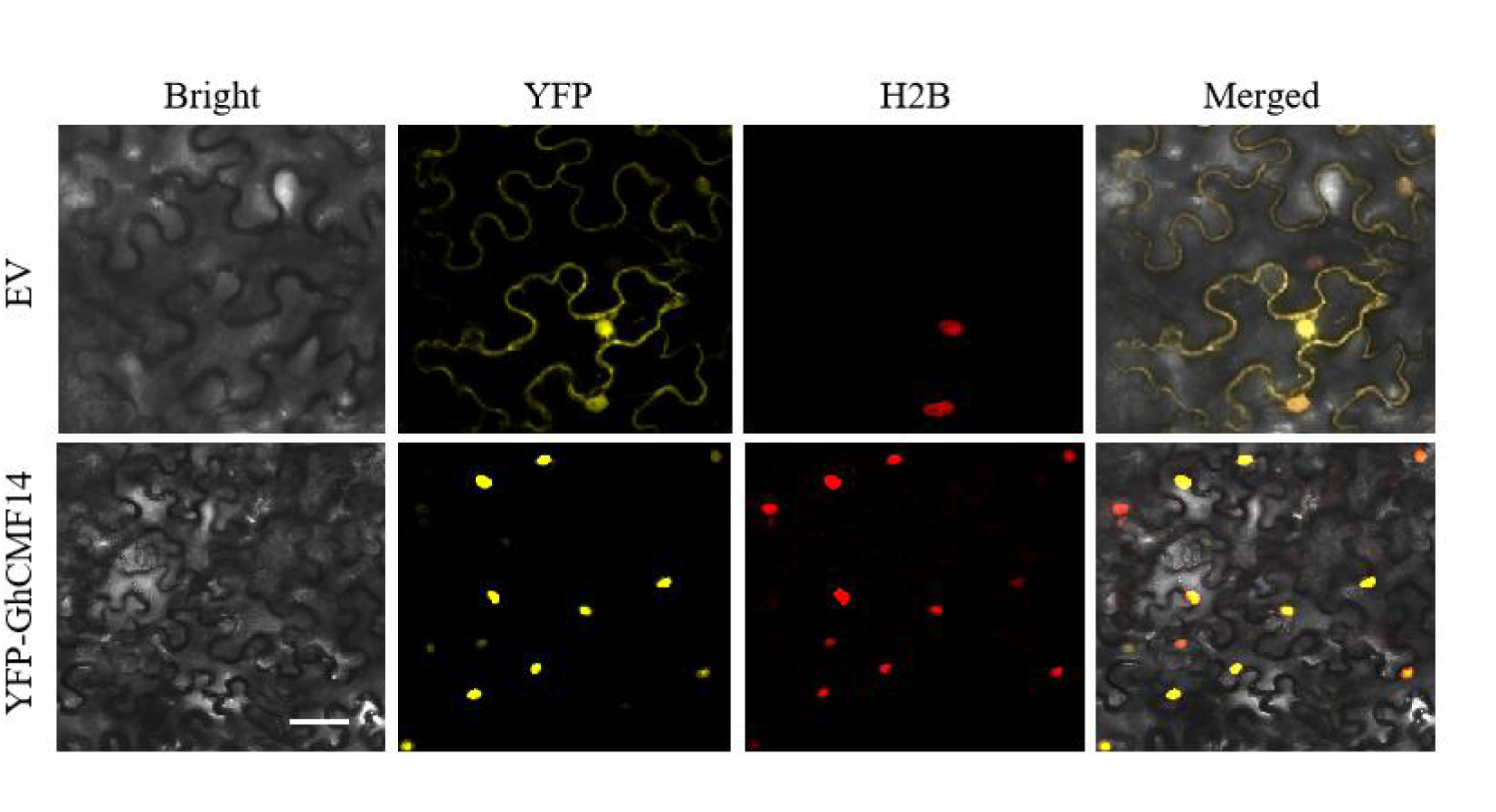
Subcellular localization of GhCMF14 in *N. bentamiana* mesophyll cells. EV: 35S-YFP empty vector; YFP-GhCMF14: localization of GhCMF14 protein; YFP: yellow fluorescence; Bright: bright field; H2B: nuclear marker H2B-mCherry ; Merge: yellow fluorescence fused with H2B and bright field. Bars = 40 μm.

### 3.9. Silencing of GhCMF14 reduces the sensitivity to salt stress

By analyzing the GhCMF protein interaction network and the expression profile of *GhCMF* gene under salt stress, we selected *GhCMF14* (*GH_A11G0139*) for further functional validation. To investigate the role of *GhCMF14* in response to salt stress, we used the virus-induced gene silencing (VIGS) technique to silence the *GhCMF14* gene in cotton via the pYL156 vector. The silencing of the phytoene desaturase (PDS) gene disrupted the carotenoid biosynthesis pathway, resulting in the loss of photoprotection and causing a bleaching effect in plants [78]. After 2 weeks infiltration injection, the leaves of pYL156: PDS plants displayed a distinct bleaching phenotype (Fig. 8A), indicating the VIGS system was successfully constructed. After 3 weeks of infiltration, we extracted total RNA from the leaves of GhCMF14-silenced plants and pYL156 plants. qRT-PCR analysis showed that the expression level of *GhCMF14* in GhCMF14-silenced plants was approximately 72% lower than that in pYL156 plants (Fig. 8B), demonstrating effective VIGS silencing. Under 300 mM NaCl treatment, the leaves of pYL156 seedlings exhibited wilting, whereas GhCMF14-silenced plants maintained normal leaf appearance (Fig. 8C), the chlorophyll content in the leaves of GhCMF14-silenced plants was markedly increased (Fig. 8D), the proline content was significantly higher than in pYL156 (Fig. 8E), and MDA content slightly decreased (Fig. 8F). These results suggested that GhCMF14-silenced plants could enhance their adaptability to salt stress by regulating osmotic pressure and photosynthesis. Research indicates that under various abiotic stresses, plants reduce damage by increasing the activity of antioxidant enzymes to eliminate excess ROS and prevent lipid peroxidation [79]. The result showed that the activities of SOD and POD in the leaves of GhCMF14-silenced plants were higher than those in pYL156 plants (Fig. 8G, 8H). We stained cotton leaves with DAB to detect H_2_O_2_ content under normal or treatment conditions in pYL156 and GhCMF14-silenced plants. DAB staining showed that the color depth of GhCMF14-silenced plants was slightly lower than that of the pYL156 plants under normal conditions. And the color depth of GhCMF14-silenced plants was substantially lower than that of the pYL156 plants under 300 mM NaCl treatment (Fig. 8I), indicating that salt treatment caused an increase in H_2_O_2_ content in pYL156 leaves, while a decrease in H_2_O_2_ content in GhCMF14-silence plant. These results indicated that lipid peroxidation and oxidative stress were lower in GhCMF14-silenced plants, and that those plants were resistant to NaCl stress by enhancing the antioxidant system to eliminate ROS accumulation, which also indicated that GhCMF14 played a negative regulatory role in salt tolerance of cotton.

**Fig. 8.**
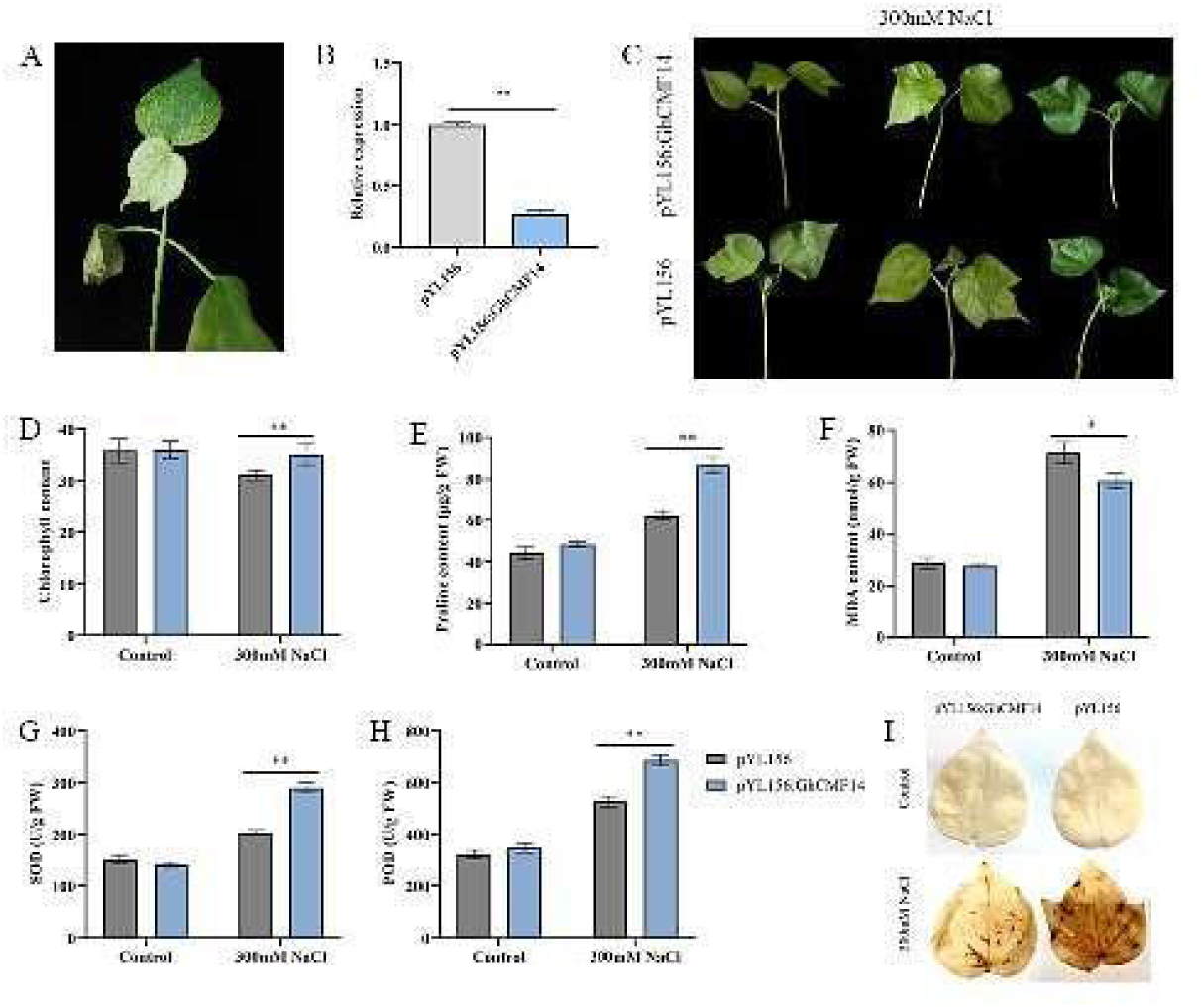
VIGS experiments of the *GhCMF14* gene and measurement of physiological indicators. (A) The true leaves of pYL156: PDS cotton showed bleaching. (B) qRT-PCR analysis of pYL156 and pYL156:GhCMF14 plants. (C) Phenotypes of pYL156 and pYL156:GhCMF14 plants under salt stress. (D-H) Physiological indicators including chlorophyII, Proline and MDA content, SOD, POD activities of pYL156 and pYL156:GhCMF14 plants. (I) DAB staining of the leaves of pYL156 and pYL156:GhCMF14 plants exposed to 300 mM NaCl for 24 h.

## 4. Discussion

In recent years, CMF proteins have attracted more and more attention due to their diverse functions in regulating plant growth and development, adjusting flowering time and coping with abiotic stress. At present, *CMF* genes has been extensively analyzed in various plant species, such as soybean, rice, poplar, and wheat. Nevertheless, studies in cotton have mainly focused on the *COL* subfamily with BBOX and CCT domain, and comprehensive analysis of *CMF* genes has not been carried out. In order to further explore the potential function of *CMF* genes in cotton, a systematic analysis of *CMF* genes in four cotton species was conducted in this study, aiming to provide more information on their evolution and function.

### 4.1. Phylogenetic analysis, conserved motifs and function of cotton CMF gene family

In this study, we identified 16, 17, 34, and 34 *CMF* genes from *G. raimondii*, *G. arboreum*, *G. hirsutum*, and *G. barbadense*, respectively (Table S1). By constructing an evolutionary tree, these *CMF* genes were divided into six distinct groups (Figure 1). Compared with *AtCMFs*, the cotton *CMFs* are more distantly related to *OsCMFs*, which is consistent with the evolutionary relationship of cotton, *Arabidopsis* and rice. The distribution of gene structure and motifs can be used as valid evidence of evolutionary relationships between species or genes [80]. Typically, members in the same group have similar exon/intron structures and motif distribution patterns, suggesting that they may have similar functions [81]. As shown in Fig. S2, similar gene structure and motif distribution in the same group provide additional evidence to validate the classification based on phylogenetic analysis. In Group VI, *FITNESS* (*AtCMF3*) plays a key role in regulating reactive oxygen species levels and defense responses [18]. *CIA2* (*AtCMF14*) and *CIL* (*AtCMF9*) coordinate chloroplast biogenesis and function mainly by up-regulating the expression of nuclear factor GOLDEN2-LIKE 1 (GLK1) and chloroplast transcription-, translation-, protein import-, and photosynthesis-related genes [19]. *OsCMF12* (*Ghd7*, *Os07g15770*) is involved in the growth and development of rice and its response to drought stress [13], thus, their corresponding homologous *GhCMF10* and *GhCMF27* may have a similar function. Of course, in order to deeply understand the functional genomics of plants, apart from referencing the homologous genes, the identification and characterization of the individual *GhCMF* gene are also essential. In addition, our study showed that cotton *CMF* genes are unevenly distributed across the chromosomes of the four cotton species (Fig. 2), which may be due to differential rates of genomic evolution and inter-genomic transmission of genetic information [82, 83].

The functional differentiation of duplicated gene pairs is the source of generating new genes in plants, which injects new impetus into the evolution of plant genome. Changing gene expression patterns is another key factor in functional differentiation [84]. Except for photoperiodic regulation, functions of *CMF* genes are also related to abiotic stress, plant hormone responsiveness, and growth regulation. The cis-acting elements contained in each gene are different. Some genes were found to have fewer cis-acting elements than others. For example, *GhCMF19* and *GhCMF29* have fewer cis-acting elements, both of them only harbor one light responsive element and one anaerobic induction responsive element. Previous studies have also reported that some genes in other gene families, such as the NFYA and AAO gene families [85, 86], have fewer cis-acting elements. By comparing cis-acting elements of *GhCMF* genes with heat maps of differential gene expression under cold, hot, NaCl and PEG stress, it was further verified that some members of CMF gene family play important roles in abiotic stress. Under different abiotic stress, *GhCMF* genes showed different expression trends. Clearly, members of *GhCMF* gene family have their own ways to regulate abiotic stress. For example, *GhCMF10*, *GhCMF14*, *GhCMF27* and *GhCMF31* genes were induced under salt stress and cold stress (Fig. 4B), and the role of these four *GhCMF* genes was further verified by analyzing their expression under NaCl treatment (Fig. 5B). Whole-genome expression profiles provide critical data needed to build co-expression networks, thus allowed us to better identify and understand biological processes and gene functions [87]. The co-expression network reflects the correlation of different gene expression patterns, and has suggestive effect in tracking the genes in the same pathway [88]. By constructing the co-expression network of *GhCMF14*, it was found that GhCMF14 protein interacts with ERCC1, CR88, ZML2, ZML1, PRR7, ARP6, BBX15, BBX14, BBX16, and AT1G49390 (Fig. 6). This provides additional evidence to verify the function of GhCMF14 gene in salt stress.

### 4.2. Evolution of CMF genes in Gossypium

Allotetraploid cotton (*G. hirsutum* and *G. barbadense*) was formed from the A and D genomes through natural hybridization and chromosome doubling [89]. In diploid cotton, each *CMF* gene corresponds to two homologous genes in its corresponding allotetraploid. However, GhCMF1 and GbCMF1 in the A subgenome of allotetraploid have no corresponding *CMF* gene in *G. arboreum*; Similarly, *GhCMF25* and *GbCMF25* have no corresponding *CMF* genes in *G. raimondii*, suggesting that these two genes may be lost after *G. arboreum*/*G. raimondii* differentiation. This inconsistency has also been found in a genus of different cotton varieties [49, 90]. Although the genome size of *G. arboreum* is about twice that of *G. raimondii*, the number of *CMF* genes in both cotton species is roughly equal, which may be caused by the insertion of long terminal repeat (LTR) retrotransposons in *G. arboreum* [38, 39]. The number of *CMF* genes in G*. hirsutum* or *G. barbadense* is roughly equal to that sum of *G. raimondii* and *G. arboreum*, suggesting that the *CMF* gene family has expanded over the course of evolution. Studies have shown that *G. raimondii* and *G. arboreum* underwent a *Gossypiuum*-specific whole-genome duplication (WGD) events [38, 39, 91] and that the *CMF* gene family of *G. raimondii* and *G. arboretum* may be amplified by an ancient WGD event. Thus, the expansion of the *CMF* gene family in *G. hirsutum* and *G. barbadense* may have been caused by hybridization and subsequent polyploid events. In principle, gene family expansion is mainly achieved through three mechanisms: segmental duplication, tandem duplication and retroposition [92, 93]. During evolution, except for small-scale tandem duplications, most segmental duplications contribute to the generation of new genes, thereby increasing the complexity of plant genomes [92]. Based on the theory that genes separated by 5 or fewer genes within the 100 kb region of the chromosome may be formed by tandem repeats [94], combined with the distribution of the *CMF* gene across the chromosomes of the four cotton species, we demonstrated that segmental duplication is largely responsible for the expansion of the *CMF* gene family in the four cotton species.

Gene duplication produces functional differences and is considered to be the most important factor in speciation and environmental adaptation [39]. At the same time, the ratio of non-synonymous to synonymous substitution (Ka/Ks) is an important indicator to evaluate the selection pressure faced by genes or a gene region [95]. In the course of evolution, genes must withstand the pressure of natural selection. We performed Ka/Ks ratio calculations for *CMF* duplicated pairs within and between *G. raimondii*, *G. arboreum*, *G. hirsutum*, and *G. barbadense* to assess their selection pressure (Table S4). After analyzing the 410 duplicated gene pairs in four cotton species, it was found that 400 pairs of them (97.56%) had a Ka/Ks ratio less than 1, indicating that they underwent strong purifying selection. Only 10 pairs (2.44%) of replicated pairs showed positive selection, suggesting that they may have undergone rapid evolution after duplication. Our study provides useful knowledge and evidence for the expansion of the *CMF* gene family in four cotton species. Therefore, we speculate that the function of most *CMF* duplicated gene pairs was conserved over the long course of evolution, which could eliminated deleterious loss of function mutations, and a new duplicated gene at both duplicate loci can be fixed and enhanced after purifying selection [96].

### 4.3. GhCMF14 was involved in salt stress resistance of cotton plants

When plants are subjected to abiotic stress, multiple signaling pathways are triggered, and a series of adaptive response mechanisms are stimulated [97]. Plants respond to stress through a series of cellular and physiological changes [98]. Proline accumulation enhances plant stress tolerance [99]. Salt stress produces a kind of hypertonic stress on plant cells, which affects cell membrane function [100]. We silenced *GhCMF14* gene by VIGS technique, and the tolerance of GhCMF14-silenced plants under salt stress was better than that of pYL156 plants. Specifically, the contents of chlorophyll, proline and POD and SOD activities of GhCMF14-silenced plants were significantly higher than those of pYL156 plants (Fig. 8D, E, G, H). The MDA content is used to indicate the product of lipid peroxidation, and to reflect the damage extent of plant caused by abiotic stress [101]. MDA content decreased slightly (Fig. 8F), indicating that GhCMF14-silenced plants were resistant to NaCl stress by enhancing the antioxidant system and eliminate ROS accumulation, and GhCMF14 played a negative regulatory role under salt tolerance in cotton. Studies have shown that when plants are stimulated by abiotic stresses, the balance between production and removal of ROS in vivo is disrupted, and ROS levels are rise, leading to oxidative damage in plants [102]. Plants can limit ROS accumulation by enhancing antioxidant systems and regulating the expression of defense-related genes, thereby protecting plant cell membranes from damage [103].

Although the molecular mechanism of *CMF* gene tolerance to salt stress in cotton remains unclear, based on all the experimental evidence, we proposed a functional model of *GhCMF14* gene enhancing salt tolerance in cotton (Fig. 9). In GhCMF14-silenced plants, the antioxidant system was enhanced due to decreased *GhCMF14* expression, thereby improving the plant’s tolerance to salt stress. Under salt stress, signals related to salt stress down-regulated the activation of *GhCMF14* gene, promoted the enhancement of photosynthesis, and enhanced the ability of cells to effectively eliminate ROS by increasing the activity of SOD and POD and the accumulation of proline in plants, thus enhancing the salt tolerance of cotton. This study provides important biological insights for future salt tolerated gene research. In addition, this study is helpful to further explore the biological characteristics, molecular functions and antioxidant effects of CMF gene in cotton under various abiotic stresses.

**Fig. 9.**
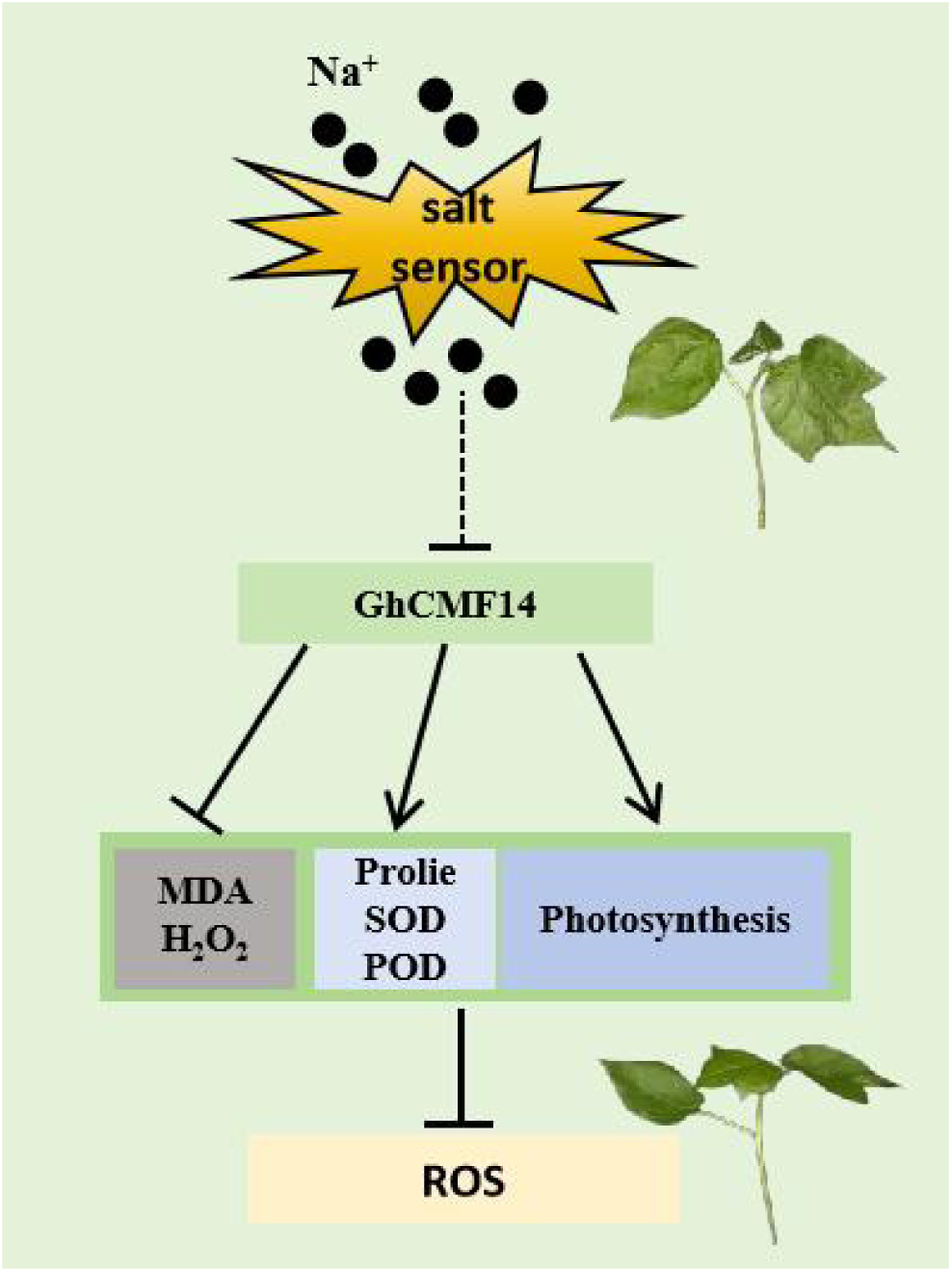
A model depicting the function of GhCMF14 in cotton for enhancing tolerance to sat stress. The solid arrows indicate activation of transcription, dotted-lines denote a relationship that has not been established as direct, and lines ending with a dash denote repression of transcription.

## 5. Conclusions

In this study, a total of 101 *CMF* genes were identified in four cotton species. The analysis of gene structure and motif composition suggested that the cotton *CMF* genes were relatively conserved. Collinearity analysis suggested that segmental duplication was the main driving force of *CMF* gene families in four cotton varieties. The expression levels of *GhCMF* genes were different in different tissues, and some were induced by various abiotic stresses. In addition, co-expression network of *GhCMF14* indicated that it interacts with other proteins to against abiotic stress. Silencing the *GhCMF14* gene, which encoded protein is localized in the nucleus, improves salt tolerance. In summary, this study lays an important foundation for the improvement of stress resistance in cotton, and provides a new insight for the functional research on *CMF* gene family.

## Credit authorship contribution statement

Yupeng Cui, Xue-Rong Zhou and Wuwei Ye designed the study. Yupeng Cui, Gongyao Shi, Zhanshuai Li analyzed the data. Yupeng Cui supervised the experiments and wrote the paper. Shuai Wang and Xingyu Peng conducted the experiment. Yupeng Cui and Wuwei Ye provided financial support. Yupeng Cui, Xue-Rong Zhou, Wuwei Ye revised the manuscript.

## Figure legend

Fig. S1. The member distribution of 135 *CMF* family members from six species in different groups.

Fig. S2. Analysis of gene structures and conserved motif elements of 101 *CMF* family members in *Gossypium*.

Fig. S3. Specific information of 3 motifs. The x-axis indicates the conserved sequences of the domain. The height of each letter indicates the conservation of each residue across all proteins. The y-axis is a scale of the relative entropy, which reflects the conservation rate of each amino acid.

Fig. S4. Syntenic relationship of *CMF* duplicated genes pairs from each of the four cotton species (*G. hirsutum*, *G. barbadense*, *G. arboreum* and *G. raimondii*).

Fig. S5. Syntenic relationship of CMF duplicated genes pairs from inter genomic combinations of the four cotton species (*G. hirsutum*: Gh, *G. barbadense*: Gb, *G. arboreum*: Ga, and *G. raimondii*: Gr).

Table S1. List of *CMF* gene family in four *Gossypium* species

Table S2. The list of primers in our study.

Table S3. Expansion of *CMF* gene family followed by gene duplication

Table S4. Selection pressure analysis of the *CMF* duplicated gene pairs from four *Gossypium* species

Table S5. The cis-elements in the 2.0 kb promoter regions of *GhCMFs*.

Table S6. The list of cis-elements involved in abiotic stresses and plant hormone response in the promoter regions of *GhCMFs*.

Table S7. The expression levels of *GhCMFs* in different tissues and abotic stress.

## Supporting information

Fig. S1, S2, S3, S4, S5, Table S1, Table S2, Table S3, and will be usde for the link to the file on the preprint site.

