## Supplementary material for "Exploring the Role of Cotton *CMF* Genes in Salt Stress Tolerance: Insights from Phylogenetic, Expression, and Functional Analyses": Fig. S1, S2, S3, S4, S5, Table S1, Table S2, Table S3, and will be usde for the link to the file on the preprint site.: Table S6.docx

**Table S6. The list of cis-elements involved in abiotic stresses and plant hormone response in the promoter regions of *GhCMFs*. 0: indicated no corresponding *cis*-element.**

| **Gene name** | **Light** | | | | | **Drought** | **Low-temperature** | **Defense and Stress** | **Auxin** | **GA** | | | **Anaerobic induction** | **MeJA** | | **ABA** |
| --- | --- | --- | --- | --- | --- | --- | --- | --- | --- | --- | --- | --- | --- | --- | --- | --- |
|  | **G-box** | **Box 4** | **GT1-motif** | **MRE** | **TCT-motif** | **MBS** | **LTR** | **TC-rich** | **TGA-element** | **GARE** | **P-box** | **TATC-box** | **ARE** | **CGTCA motif** | **TGACG-motif** | **ABRE** |
| *GhCMF1* | 1 | 1 | 1 | 1 | 1 | 1 | 0 | 0 | 0 | 1 | 1 | 0 | 0 | 0 | 0 | 1 |
| *GhCMF2* | 1 | 1 | 0 | 1 | 1 | 1 | 0 | 1 | 1 | 0 | 0 | 0 | 1 | 1 | 1 | 1 |
| *GhCMF3* | 0 | 1 | 0 | 1 | 0 | 0 | 0 | 1 | 0 | 0 | 1 | 0 | 1 | 0 | 0 | 0 |
| *GhCMF4* | 1 | 1 | 1 | 1 | 1 | 0 | 0 | 1 | 1 | 0 | 0 | 0 | 1 | 0 | 0 | 1 |
| *GhCMF5* | 1 | 1 | 1 | 0 | 1 | 0 | 0 | 0 | 0 | 0 | 0 | 1 | 1 | 1 | 1 | 1 |
| *GhCMF6* | 1 | 1 | 0 | 0 | 1 | 0 | 0 | 0 | 1 | 1 | 0 | 0 | 1 | 1 | 1 | 1 |
| *GhCMF7* | 1 | 1 | 0 | 0 | 1 | 1 | 0 | 1 | 1 | 0 | 0 | 0 | 1 | 1 | 0 | 1 |
| *GhCMF8* | 1 | 1 | 1 | 1 | 1 | 1 | 0 | 1 | 1 | 0 | 0 | 0 | 1 | 1 | 1 | 1 |
| *GhCMF9* | 1 | 0 | 1 | 0 | 0 | 1 | 0 | 0 | 0 | 0 | 1 | 0 | 1 | 1 | 1 | 1 |
| *GhCMF10* | 1 | 1 | 0 | 0 | 1 | 0 | 1 | 1 | 0 | 0 | 0 | 0 | 1 | 1 | 1 | 1 |
| *GhCMF11* | 0 | 1 | 1 | 1 | 1 | 0 | 0 | 1 | 0 | 0 | 0 | 0 | 1 | 0 | 0 | 0 |
| *GhCMF12* | 0 | 1 | 1 | 0 | 0 | 1 | 1 | 0 | 1 | 0 | 0 | 0 | 1 | 1 | 1 | 0 |
| *GhCMF13* | 1 | 1 | 1 | 0 | 1 | 0 | 0 | 0 | 1 | 0 | 0 | 0 | 0 | 1 | 1 | 1 |
| *GhCMF14* | 1 | 1 | 1 | 1 | 1 | 1 | 0 | 0 | 1 | 0 | 0 | 0 | 1 | 1 | 1 | 0 |
| *GhCMF15* | 1 | 1 | 1 | 0 | 1 | 0 | 0 | 0 | 0 | 0 | 0 | 0 | 0 | 1 | 1 | 1 |
| *GhCMF16* | 1 | 1 | 0 | 0 | 0 | 0 | 0 | 0 | 0 | 0 | 1 | 1 | 1 | 1 | 1 | 1 |
| *GhCMF17* | 1 | 1 | 0 | 0 | 0 | 0 | 0 | 0 | 1 | 0 | 1 | 0 | 1 | 1 | 1 | 1 |
| *GhCMF18* | 1 | 1 | 0 | 0 | 1 | 0 | 1 | 0 | 1 | 0 | 1 | 0 | 0 | 0 | 0 | 1 |
| *GhCMF19* | 0 | 0 | 1 | 0 | 0 | 0 | 0 | 0 | 0 | 0 | 0 | 0 | 1 | 0 | 0 | 0 |
| *GhCMF20* | 1 | 1 | 0 | 0 | 1 | 0 | 1 | 1 | 1 | 0 | 1 | 0 | 0 | 0 | 1 | 1 |
| *GhCMF21* | 1 | 1 | 1 | 0 | 1 | 1 | 1 | 1 | 0 | 0 | 1 | 0 | 1 | 0 | 0 | 1 |
| *GhCMF22* | 1 | 1 | 1 | 0 | 0 | 0 | 1 | 1 | 1 | 0 | 0 | 1 | 1 | 1 | 1 | 1 |
| *GhCMF23* | 0 | 1 | 0 | 0 | 0 | 0 | 0 | 0 | 1 | 1 | 1 | 0 | 1 | 1 | 1 | 0 |
| *GhCMF24* | 1 | 1 | 1 | 0 | 0 | 1 | 1 | 0 | 1 | 0 | 1 | 0 | 1 | 1 | 1 | 0 |
| *GhCMF25* | 1 | 1 | 0 | 1 | 1 | 1 | 1 | 1 | 0 | 0 | 0 | 0 | 0 | 1 | 1 | 1 |
| *GhCMF26* | 1 | 1 | 1 | 0 | 0 | 1 | 1 | 0 | 0 | 0 | 0 | 0 | 1 | 1 | 1 | 1 |
| *GhCMF27* | 0 | 1 | 1 | 1 | 0 | 0 | 0 | 0 | 1 | 0 | 0 | 0 | 1 | 1 | 1 | 0 |
| *GhCMF28* | 0 | 1 | 0 | 0 | 1 | 0 | 0 | 0 | 1 | 0 | 1 | 0 | 1 | 1 | 1 | 0 |
| *GhCMF29* | 0 | 0 | 1 | 0 | 0 | 0 | 0 | 0 | 0 | 0 | 0 | 0 | 1 | 0 | 0 | 0 |
| *GhCMF30* | 1 | 1 | 1 | 1 | 1 | 1 | 0 | 1 | 1 | 0 | 0 | 1 | 1 | 1 | 1 | 0 |
| *GhCMF31* | 0 | 1 | 1 | 1 | 0 | 0 | 0 | 0 | 1 | 0 | 1 | 0 | 1 | 1 | 1 | 0 |
| *GhCMF32* | 0 | 1 | 1 | 0 | 1 | 0 | 1 | 0 | 0 | 0 | 1 | 0 | 0 | 1 | 1 | 1 |
| *GhCMF33* | 0 | 1 | 1 | 0 | 0 | 0 | 0 | 0 | 0 | 0 | 1 | 0 | 1 | 0 | 0 | 0 |
| *GhCMF34* | 1 | 1 | 1 | 0 | 0 | 1 | 0 | 1 | 1 | 0 | 0 | 1 | 1 | 1 | 1 | 1 |
