## Supplementary figures and images for "Exploring the Role of Cotton *CMF* Genes in Salt Stress Tolerance: Insights from Phylogenetic, Expression, and Functional Analyses"

### Fig. S1.tif

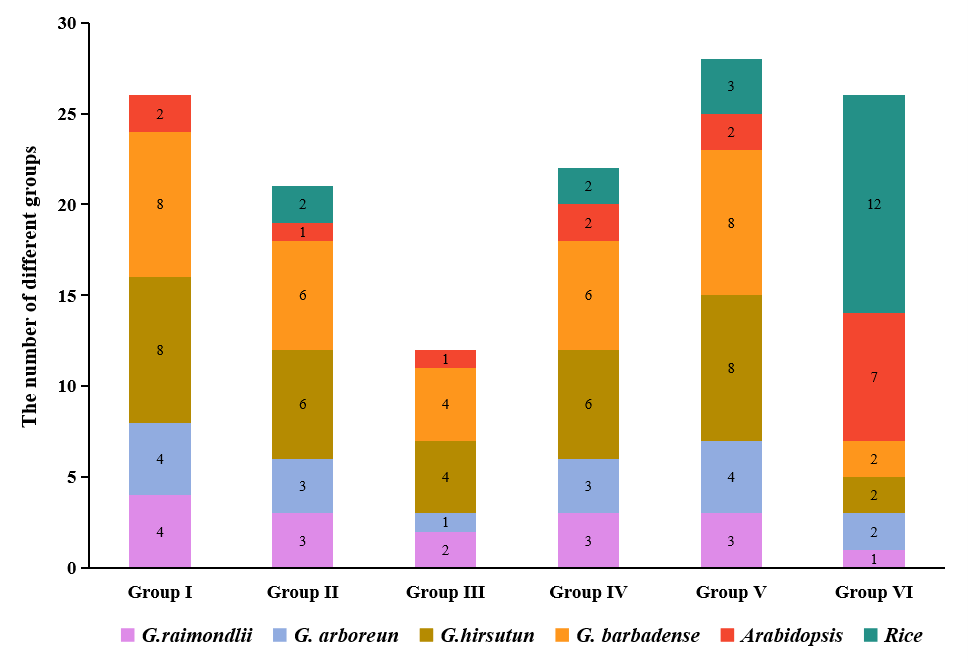

### Fig. S2.tif

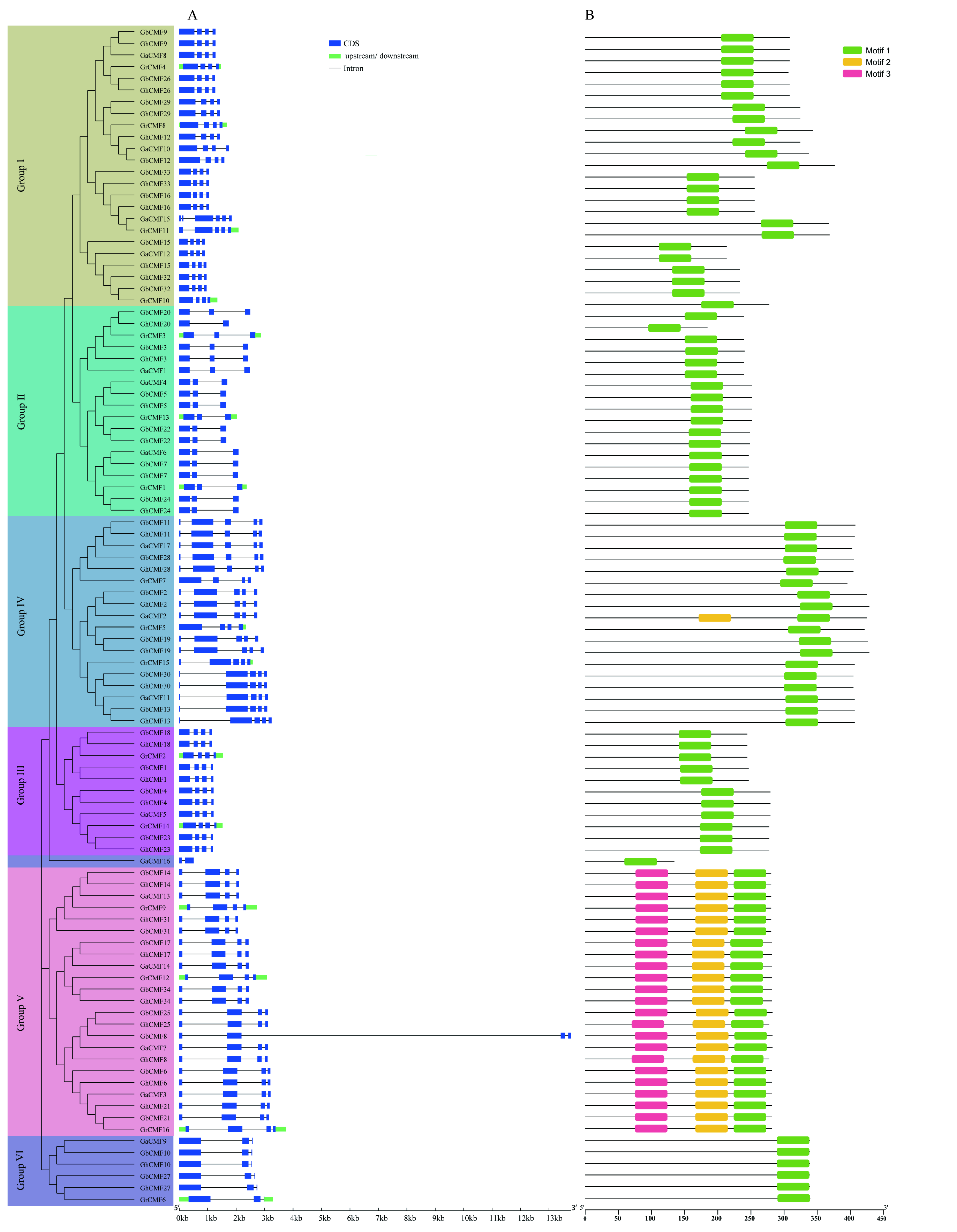

### Fig. S3.tif

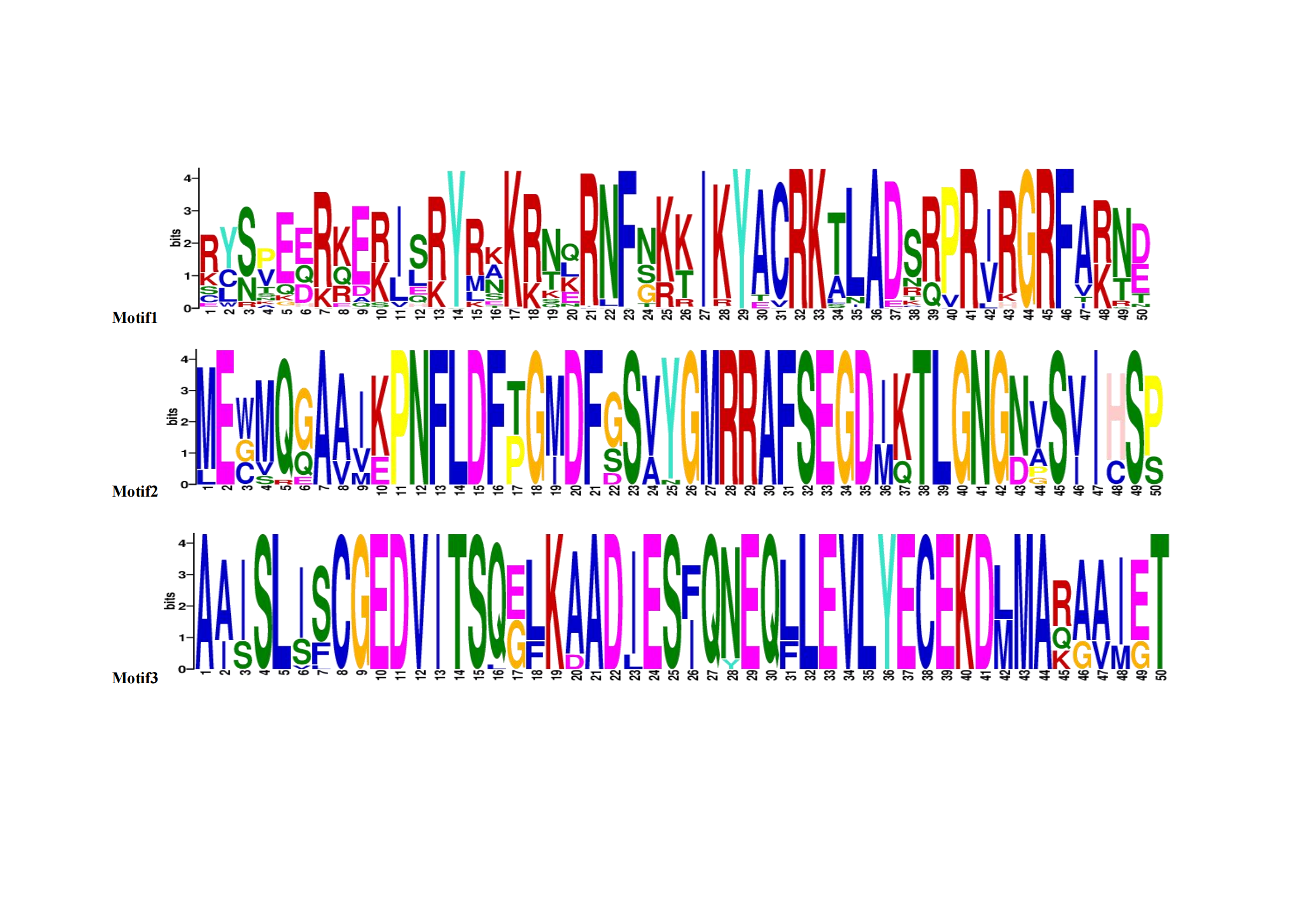

### Fig. S4.tif

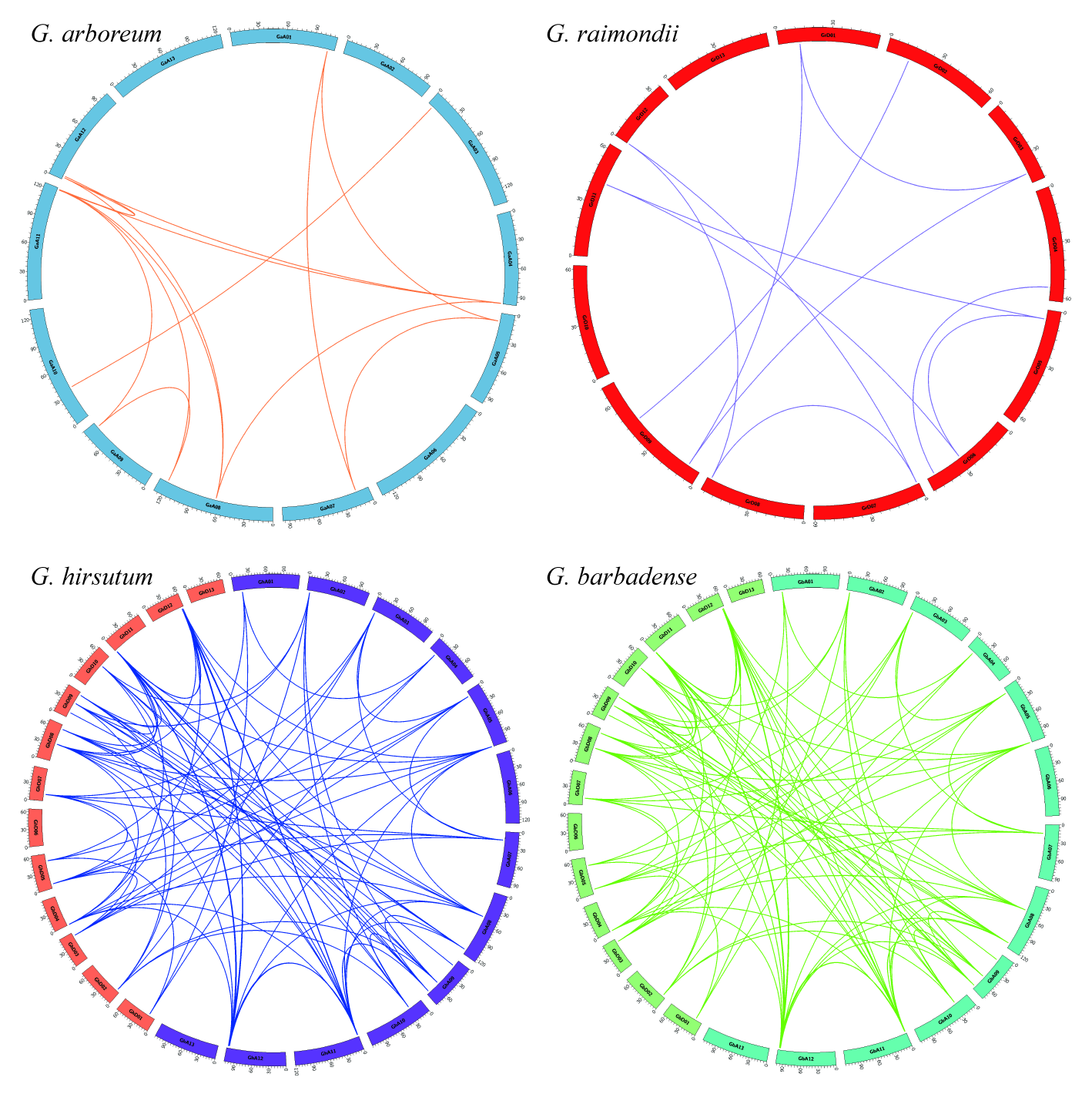

### Fig. S5.tif

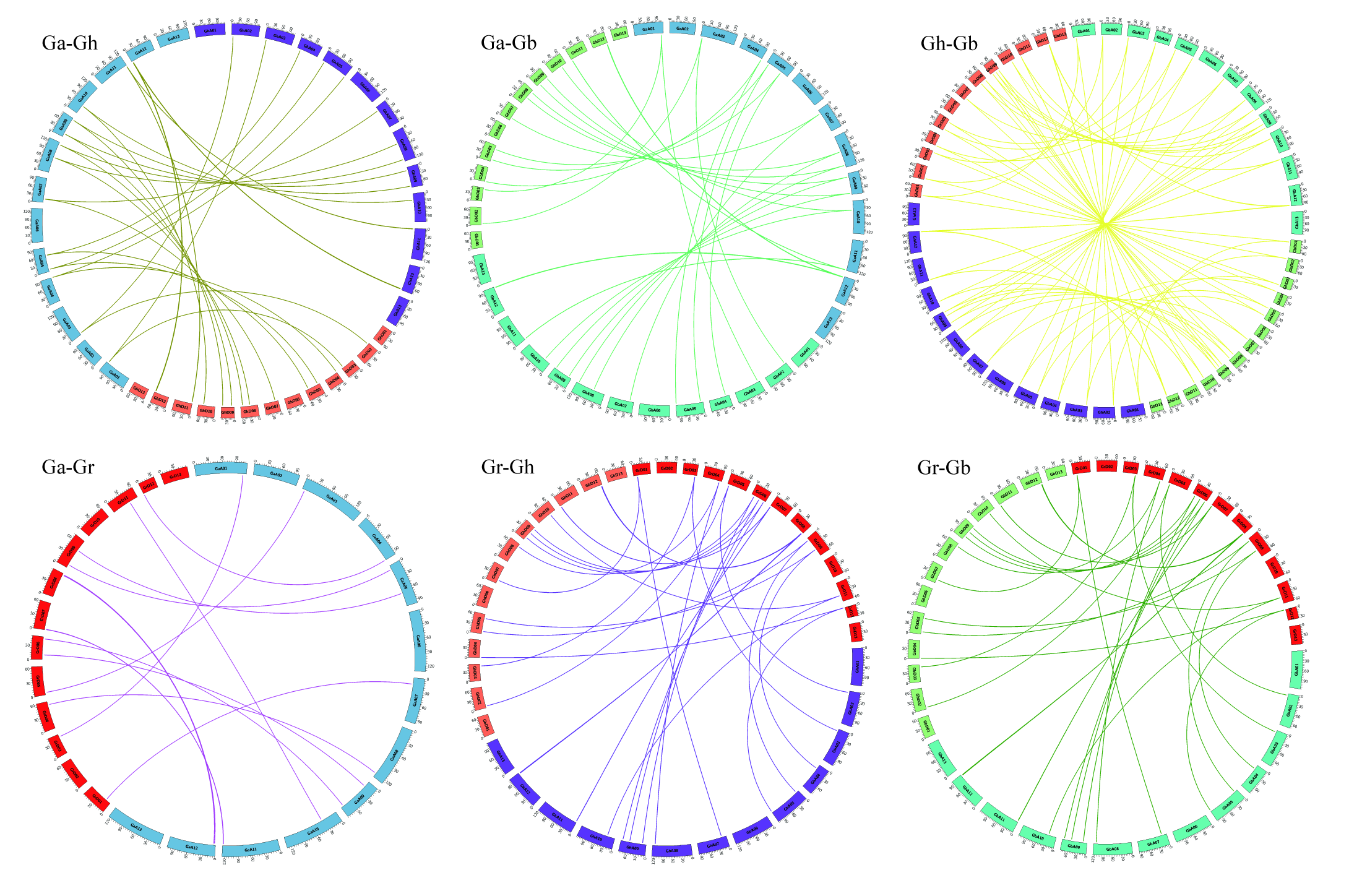
